# The 18S rRNA Methyltransferase DIMT-1 Regulates Lifespan in the Germline Later in Life

**DOI:** 10.1101/2024.05.14.594211

**Authors:** M. Hafiz Rothi, Gautam Chandra Sarkar, Joseph Al Haddad, Wayne Mitchell, Kejun Ying, Nancy Pohl, Roberto G. Sotomayor-Mena, Julia Natale, Scarlett Dellacono, Vadim N. Gladyshev, Eric Lieberman Greer

**Author notes:** Correspondence: Eric L. Greer. These authors contributed equally to this work (M. Hafiz Rothi, Gautam Chandra Sarkar, Joseph Al Haddad).

## Abstract

Ribosome heterogeneity has emerged as an important regulatory control feature for determining which proteins are synthesized, however, the influence of age on ribosome heterogeneity is not fully understood. Whether mRNA transcripts are selectively translated in young versus old cells and whether dysregulation of this process drives organismal aging is unknown. Here we examined the role of ribosomal RNA (rRNA) methylation in maintaining appropriate translation as organisms age. In a directed RNAi screen, we identified the 18S rRNA N6’-dimethyl adenosine (m^6,2^A) methyltransferase, *dimt-1,* as a regulator of *C. elegans* lifespan and stress resistance. Lifespan extension induced by *dimt-1* deficiency required a functional germline and was dependent on the known regulator of protein translation, the Rag GTPase, *raga- 1,* which links amino acid sensing to the mechanistic target of rapamycin complex (mTORC)1. Using an auxin-inducible degron tagged version of *dimt-1,* we demonstrate that DIMT-1 functions in the germline after mid-life to regulate lifespan. We further found that knock-down of *dimt-1* leads to selective translation of transcripts important for stress resistance and lifespan regulation in the *C. elegans* germline in mid-life including the cytochrome P450 *daf-9,* which synthesizes a steroid that signals from the germline to the soma to regulate lifespan. We found that *dimt-1* induced lifespan extension was dependent on the *daf-9* signaling pathway. This finding reveals a new layer of proteome dysfunction, beyond protein synthesis and degradation, as an important regulator of aging. Our findings highlight a new role for ribosome heterogeneity, and specific rRNA modifications, in maintaining appropriate translation later in life to promote healthy aging.

## *dimt-1* rRNA methyltransferase knock-down extends lifespan and increases resistance to stress

Previous genome-wide RNAi screens for genes that regulate lifespan in *C. elegans* have been performed in worms where progeny production was inhibited^1–3^, which has been shown to mask the effect of some genes on lifespan^4,5^. We performed a targeted RNAi screen for putative rRNA methyltransferases present in fertile *C. elegans* to test whether manipulation of rRNA methylation could regulate lifespan or stress resistance. We knocked-down 26 putative rRNA methyltransferases and discovered that knock-down of several rRNA methyltransferases could significantly shorten or extend *C. elegans* lifespan (**Fig. 1a**). As previously reported^6^, knock-down of *T01C3.7/fib-1* extended *C. elegans* lifespan (∼9% p<0.0005, **Fig. 1a**). FIB-1 is a homologue of the 2’-O-methyltransferase fibrillarin, which is conserved in animals and plants. We also found that knock-down of *E02H1.1/dimt-1* caused the most significant extension of lifespan (22-33% p<0.0001) (**Fig. 1a**). We had previously reported that DIMT-1 is the 18S rRNA N6- dimethyladenosine (m^6,2^A) methyltransferase for adenosines 1735 and 1736 in *C. elegans*^7^. DIMT- 1 is a conserved methyltransferase that methylates conserved adjacent adenosines on the 18S rRNA in yeast and humans^8–10^.

**Fig. 1.**
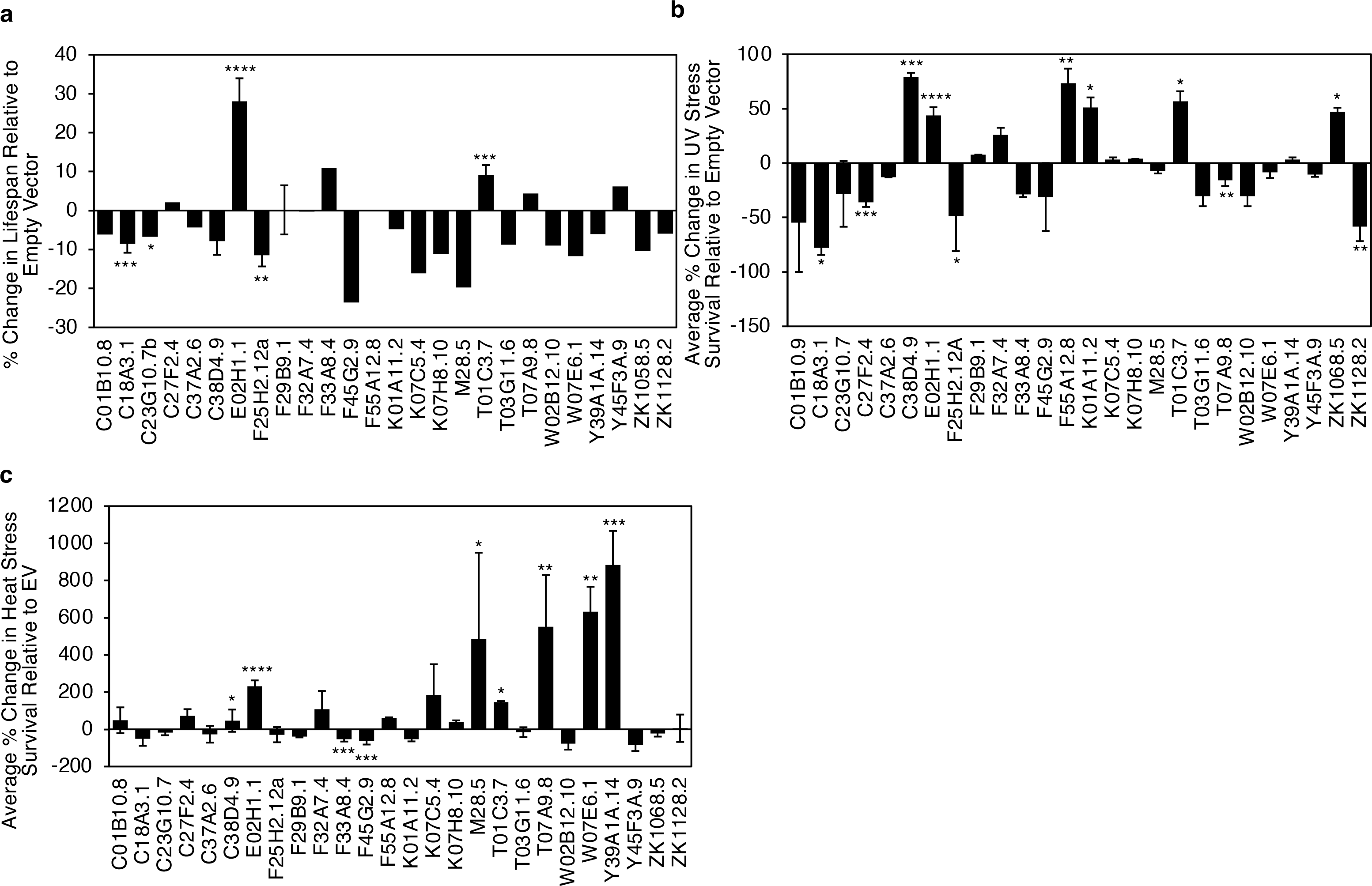
Ribosomal RNA methyltransferases regulate lifespan, heat and UV stress response. a-c, Directed RNAi screen of putative rRNA methyltransferases reveals changes in **a**) lifespan, **b**) UV stress survival, and **c**) heat stress survival relative to empty vector control worms. * p < 0.05, ** p < 0.01, *** p < 0.001, **** p < 0.0001. Some RNAi clones were not replicated (without error bars) due to no effect being observed.

Although lifespan extension can be beneficial, it may also come with detrimental effects to the organism. To determine if the effects on longevity by rRNA methyltransferase knock-down were associated with improved health, we performed UV stress and heat stress survival assays using the same targeted RNAi screen. As we had found previously^11^, knock-down of *C38D4.9/metl-5* increased both UV stress and heat stress resistance (79%, p = 0.001 and 46.5% p < 0.05, **Fig. 1b,c**). Additionally, as previously reported^12^, knock-down of *W07E6.1/nsun-1* increased heat stress resistance without having a significant effect on overall lifespan (633%, p=0.0036 and -11.7%, p =0.6713, **Fig. 1**). *E02H1.1/dimt-1* also displayed a significant increase in both UV stress (43.7% p<0.0001) and heat stress resistance (231.3% p<0.0001) (**Figs. 1b, c**). As *dimt-1* knock-down caused the most significant lifespan extension (**Fig. 1a**), we focused subsequent analyses on determining how DIMT-1 can regulate longevity. To determine whether *dimt-1* was regulating longevity by altering protein homeostasis, we measured the levels of 3 stress induced chaperone proteins important for stabilizing misfolded proteins, HSP-4^13,14^, HSP-6^15,16^, and HSP-16.2^17,18^. These chaperones are indicators of misfolded protein loading in response to endoplasmic reticulum (ER) stress, mitochondrial stress, and cytosolic stress, respectively. While HSP-6 and HSP-16.2 showed no change upon *dimt-1* knock-down (Extended Data Fig. 1b, c), HSP-4 had a significant reduction in expression levels when *dimt-1* was knocked down, indicating that when levels of DIMT-1 are decreased, the level of misfolded protein loading in the endoplasmic reticulum (ER) is also lowered, likely due to better protein turnover compared to control (Extended Data Fig. 1d). To determine whether there is a better ER specific protein turnover, we performed a proteotoxicity survival assay using the ER stress inducer tunicamycin (TM), which block N-linked glycosylation. We found that *dimt-1* knock-down caused a significant increase in tunicamycin survival relative to WT control worms (Extended Data Fig. 1e).

Taken together, these findings suggest that rRNA methyltransferases play significant roles in regulating lifespan and responses to UV and heat stress and more specifically that *dimt-1* deficiency increases longevity and stress resistance.

## rRNA modifications are dynamically regulated throughout the lifespan

Although we found that rRNA methyltransferases regulate lifespan and stress resistance, it was still unclear whether rRNA modifications themselves change during the life of the organism. Changes in specific modifications with age may indicate a regulatory role for rRNA methylation events. We first wanted to see if the rRNA methyltransferase genes are dynamically expressed during the life of *C. elegans.* Examination of previously published transcriptional profiles of *C. elegans* during aging^19^, revealed that a number of putative rRNA methyltransferases are dynamically expressed with age (Extended Data Fig. 2a). There were genes which are strongly expressed in early life and declined towards the end such as *ZK1128.2/mett-10, W01B11.3/nol-58,* and *W07E6.1/nsun-1*. Alternatively, genes such as *E02H1.1/dimt-1* and *C18A3.1/damt-1* showed the opposite trend, peaking near the end of the lifespan (Extended Data Fig. 2a). We independently confirmed that *E02H1.1/dimt-1* expression increases as *C. elegans* age (Extended Data Fig. 2b) suggesting that knock-down of *dimt-1* is reverting the worm to a more youthful state.

To test if the rRNA modifications themselves showed similar patterns during aging, we performed ultra-high performance liquid chromatography coupled with mass spectrometry (UHPLC-MS) with wildtype (WT) N2 worms harvested in an age gradient at days 4, 7, 10, 13, 16, and 19. We measured rRNA modification levels in the 26S and 18S rRNA subunits in 4 biological replicates. We observed a similarly dynamic pattern of change in rRNA modifications across lifespan (**Fig. 2a**). 2’-O-ribose methylation increased on all four nucleosides (Am, Cm, Gm, Um) as the worm gets older in both the 26S and 18S rRNAs (**Fig. 2a** and Extended Data Fig. 2c). Other 18S rRNA modifications such as m^6^A and surprisingly, m^6,2^A, showed the inverse trend with a higher level in early life which declines as the worm ages (**Fig. 2a** and Extended Data Fig. 2c). Since knock-down of the 18S rRNA m^6,2^A methyltransferase, *dimt-1,* increased *C. elegans* lifespan we had hypothesized that m^6,2^A would have increased as *C. elegans* age. This finding could suggest a dynamic and tissue specific change in 18S rRNA m^6,2^A which might be masked by examination of changes in rRNA methylation across all tissues. These results suggest that both rRNA methyltransferases and modifications are dynamically regulated throughout the lifespan of *C. elegans*.

**Fig. 2.**
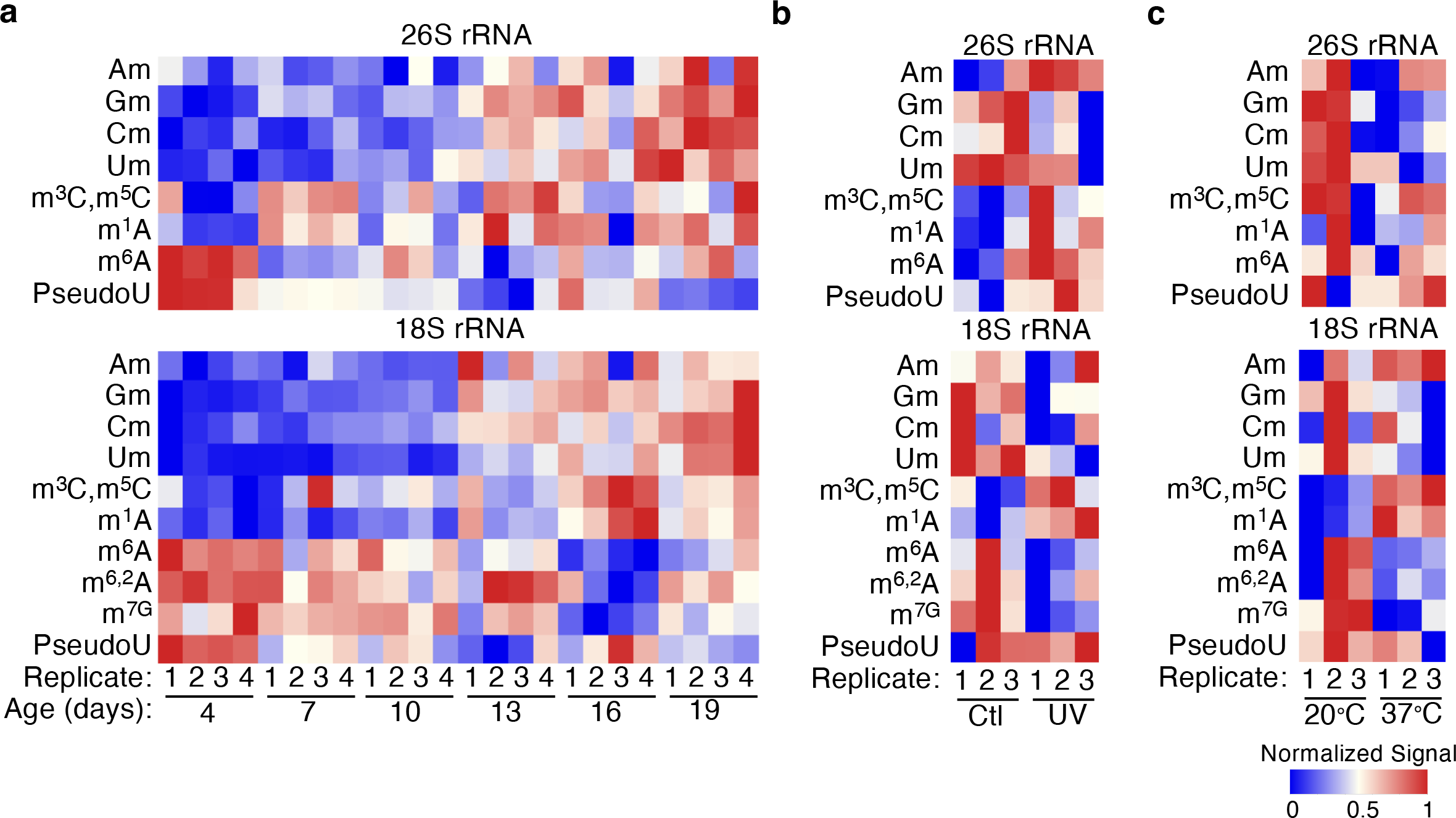
Ribosomal RNA modifications in 26S and 18S are dynamically regulated throughout life and change in response to UV and heat stress. a, 26S and 18S rRNA extracted from *C. elegans* at different ages from 4 days to 19 days reveal changes in rRNA methylation as assessed by UHPLC-ms/ms. This heat map represents the relative changes of methylation in 4 biological replicates. Changes in individual modifications can be seen in fig. S2C. **b-c**, 26S and 18S rRNA extracted from *C. elegans* after UV stress (b) or 37°C heat stress (c) relative to worms not exposed to UV (ctl) or grown at 20°C reveal some changes in rRNA methylation as assessed by UHPLC-ms/ms. These heat maps represent the relative changes of methylation in 3 biological replicates. Changes in individual modifications can be seen in Extended Data Fig. 2e-f.

We had found that the lifespan extension phenotypes observed from the RNAi screen is sometimes associated with increased stress resistance in survival assays **(Fig. 1**). We wanted to determine if there were associated changes in rRNA methyltransferases and modifications when *C. elegans* are exposed to stress. When we examined a previously published transcriptomic dataset^11^, we found that most rRNA methyltransferases increased expression in response to 37°C heat stress (Extended Data Fig. 2d). We found that rRNA modifications in wildtype N2 worms were less consistently dynamic in response to 37°C heat shock or UV stress exposure (**Fig. 2b,c**). Some rRNA modifications did change in response to the stresses, for example, 2’-O-ribose adenosine methylation (Am) in 26S rRNA was significantly increased and 2’-O-ribose cytosine methylation (Cm) in 18S rRNA subunit was significantly decreased in response to heat stress (Extended Data Fig. 2e). We also observed a significant increase in 2’-O-ribose guanosine methylation (Gm) and Am in 26S rRNA in response to UV exposure (Extended Data Fig. 2f). As with the changes in rRNA methylation during aging (**Fig. 2a**), lack of changes in specific rRNA methylation in response to stress could reflect tissue specific changes, which change in opposite directions in different tissues or an inability to capture the correct window to observe changes due to dynamic rRNA modification changes after stress exposure. Nevertheless, taken together, these results indicate that rRNA methyltransferases and rRNA modifications are dynamically regulated during the lifespan of *C. elegans*, and can undergo changes in response to environmental stresses. This suggests that rRNA modifications could be the regulatory changes that rRNA methyltransferases induce to regulate longevity and stress resistance.

## dimt-1 catalytic activity is required for lifespan and stress resistance regulation

Due to the paradoxical decrease in m^6,2^A in whole worms as *C. elegans* age when depletion of *dimt-1,* the m^6,2^A methyltransferase^7^, causes an extension in organismal lifespan we wished to determine whether DIMT-1’s regulation of lifespan and stress resistance was dependent on its catalytic activity. We generated a catalytic inactive DIMT-1 by mutating the conserved glutamic acid E79 to alanine that we had previously demonstrated was necessary for DIMT-1’s catalytic activity *in vitro*^7^. We found that *dimt-1* (E79A) mutant worms were viable but had no detectable 18S rRNA m^6,2^A (**Fig. 3a**). *dimt-1* (E79A) mutant worms displayed a significant increase in lifespan (39.49% p<0.0001, **Fig. 3b**) and a significant increase in response to the ER stress inducer tunicamycin (Extended Data Fig. 1e). Therefore, DIMT-1 methyltransferase activity is required for lifespan and stress resistance regulation.

**Fig. 3.**
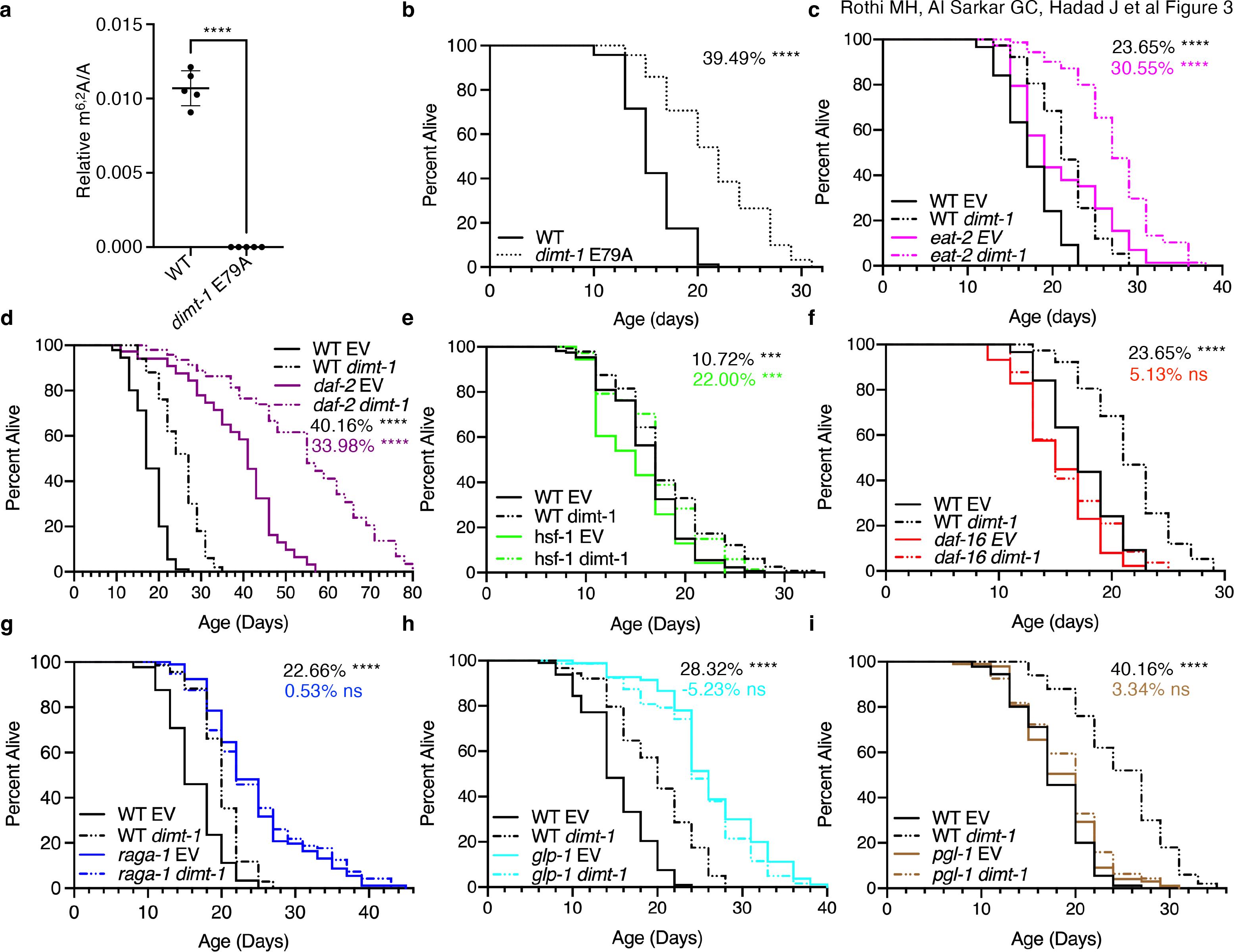
Longevity induced by *dimt-1* deficiency requires the FoxO and TOR signaling pathways and requires an intact germline. a, Mutation of glutamic acid 79 to an alanine (E79A) in *dimt-1* caused a complete elimination of 18S rRNA m^6,2^A as assessed by UHPLC-ms/ms. Statistics represent an unpaired t-test with Welch’s correction. **b,** Mutation of E79A in *dimt-1* caused lifespan extension relative to WT worms. **c,** *dimt-1* knock-down extends the lifespan of both WT and *eat-2(ad1116)* mutant worm lifespan to a similar extent (p=0.7021 by 2-way ANOVA). **d,** *dimt-1* knock-down extends the lifespan of both WT and *daf-2(e1370)* mutant worm lifespan to a similar extent (p=0.0806 by 2-way ANOVA). **e**, *dimt-1* knock-down extends the lifespan of both WT and *hsf-1(sy441)* mutant worm lifespan to a similar extent (p=0.4245 by 2- way ANOVA). **f,** *dimt-1* knock-down extends the lifespan of WT but not *daf-16(mu86)* mutant worm lifespan (p<0.0001 by 2-way ANOVA). **g,** *dimt-1* knock-down extends the lifespan of WT but does not further extend the long lifespan of *raga-1(ok386)* mutant worms (p=0.0327 by 2-way ANOVA). **h,** *dimt-1* knock-down extends the lifespan of WT but does not further extend the long lifespan of germline deficient *glp-1(e2141ts)* mutant worms that were shifted to the restrictive temperature at the L1 stage (p<0.0001 by 2-way ANOVA). **i,** *dimt-1* knock-down extends the lifespan of WT but not sterile *pgl-1(bn101ts)* mutant worms whose mothers were shifted to the restrictive temperature (25.5°C) at the L4 stage (p<0.0001 by 2-way ANOVA). Statistics and replicate experiments are presented in Supplementary Table 1. ns; not-significant, *; p < 0.05, **; p < 0.01, ***; p < 0.001, ****; p < 0.0001 as calculated by log-rank (mantel-cox) statistical test.

## *dimt-1* regulates lifespan through the DAF-16/FOXO and TOR pathways and requires the germline

Previous studies have elucidated various regulators of lifespan which are involved in pathways such as insulin-signaling, heat shock response, and target of rapamycin (TOR)^20,21^. In order to identify putative mechanisms by which DIMT-1 could regulate lifespan, we performed genetic epistasis experiments by measuring lifespans of wildtype worms and mutants of specific longevity pathways, grown on either empty vector control or *dimt-1* RNAi plates. We found that *dimt-1* knock-down increased lifespan to a similar extent in mutants of the acetylcholine receptor, *eat-2,* which has reduced pharyngeal pumping and therefore decreased food intake, and has been proposed as a genetic mimic of dietary restriction^22,23^ (23.65% in WT, 30.55% in *eat-2*; p=0.7021 by 2-way ANOVA, **Fig. 3c**). Similarly, *dimt-1* knock-down increased lifespan to a similar extent in wildtype worms and in mutants of the insulin receptor, *daf-2*^24^ (40.16% in WT, 33.98% in daf- 2; p= 0.0806 by 2-way ANOVA, **Fig. 3d**), mutants of the ubiquinone biosynthesis gene, *clk-1*^23^ (23.65% in WT, 35.2% in clk-1; p=0.0855 by 2-way ANOVA, Extended Data Fig. 3a), and in mutants of the heat-shock response transcription factor, *hsf-1*^25,26^ (10.72% in WT, 22.00% in *hsf- 1*; p=0.4245 by 2-way ANOVA, **Fig. 3e**).

Next, we tested if *daf-16,* which is a FOXO transcription factor that mediates longevity regulation downstream of several signaling pathways^26–32^, has an effect on *dimt-1* knock-down-induced lifespan extension. We found that *dimt-1* knock-down extended wildtype worm lifespan but failed to increase the lifespan of *daf-16* mutant *C. elegans* (p=0.0907), indicating that *daf-16* is functioning in the same genetic signaling pathway as *dimt-1* (23.65% in WT, 5.13% in *daf-16*; p<0.0001 by 2-way ANOVA, **Fig. 3f**). We also tested *raga-1*, which is a Rag GTPase that links amino acid sensing to mechanistic target of rapamycin complex (mTORC)1, where *raga-1* mutation causes an increase in lifespan^33^. Again *dimt-1* knock-down increased WT worm lifespan but failed to increase the long lifespan of *raga-1* mutant worms (22.67% in WT, 0.53% in raga-1, p=0.0327 by 2-way ANOVA, **Fig. 3g**), suggesting that DIMT-1 functions in the same signaling pathway as RAGA-1 and TOR. To determine if an intact germline is necessary for *dimt-1* induced lifespan extension, we performed *dimt-1* knock-down in *glp-1(e2141ts)* mutant worms, which develop 5-15 meiotic germ cells instead of ∼1,500 when shifted to the restrictive temperature^34^ and *pgl-1(bn101ts)* mutant worms, which have defective germ granules and are sterile^35^. Knock- down of *dimt-1* extended the lifespan of WT worms but failed to increase the lifespan of either *glp-1(e2141ts)* or *pgl-1(bn101ts)* mutant worms (28.32% or 40.16% in WT, -5.23% in *glp-1,* 3.34% in *pgl-1*; p<0.0001 by 2-way ANOVA, **Fig. 3h, i**), suggesting that a functional germline is required for the *dimt-1* knock-down induced lifespan extension. Taken together, these results show that *dimt-1* is likely affecting or regulating lifespan through the DAF-16 and TOR pathways and either functions in or requires the germline for longevity regulation.

## *dimt-1* functions in the germline to regulate lifespan

To resolve the paradox of depletion of *dimt-1* extending lifespan while 18S rRNA m^6,2^A increases in whole worms with age (**Fig. 2a** and Extended Data Fig. 2c), we next wished to determine whether DIMT-1 was functioning in specific tissues to regulate lifespan. To examine DIMT-1’s tissue specific function we created an auxin-inducible degron (AID) tagged DIMT-1 worm strain and crossed it with strains that express TIR1 in a tissue-specific manner to allow for tissue specific auxin-dependent depletion of DIMT-1^36,37^. We found that DIMT-1 depletion ubiquitously led to a significant decrease in 18S rRNA m^6,2^A relative to control strains as assessed by UHPLC-MS/MS, suggesting that our AID-tagged DIMT-1 strains were effective (**Fig. 4a**). We found that when DIMT-1 was depleted using two independent ubiquitous *eft-3* promoters or the germline specific *mex-5* promoter to drive TIR1 expression we observed a significant lifespan extension relative to control AID-tagged DIMT-1 strains that did not express TIR1 (29.71% p<0.0001 and 47.97% p<0.0001 respectively, **Fig. 4b** and Extended Data Fig. 3b). In contrast, we observed no lifespan extension when DIMT-1 was depleted in muscle, intestine, or neurons (**Fig. 4b**). This finding suggests that DIMT-1 not only requires the presence of a functional germline to regulate lifespan (**Fig. 3h, i**), but indeed functions in the germline to regulate lifespan.

**Fig. 4.**
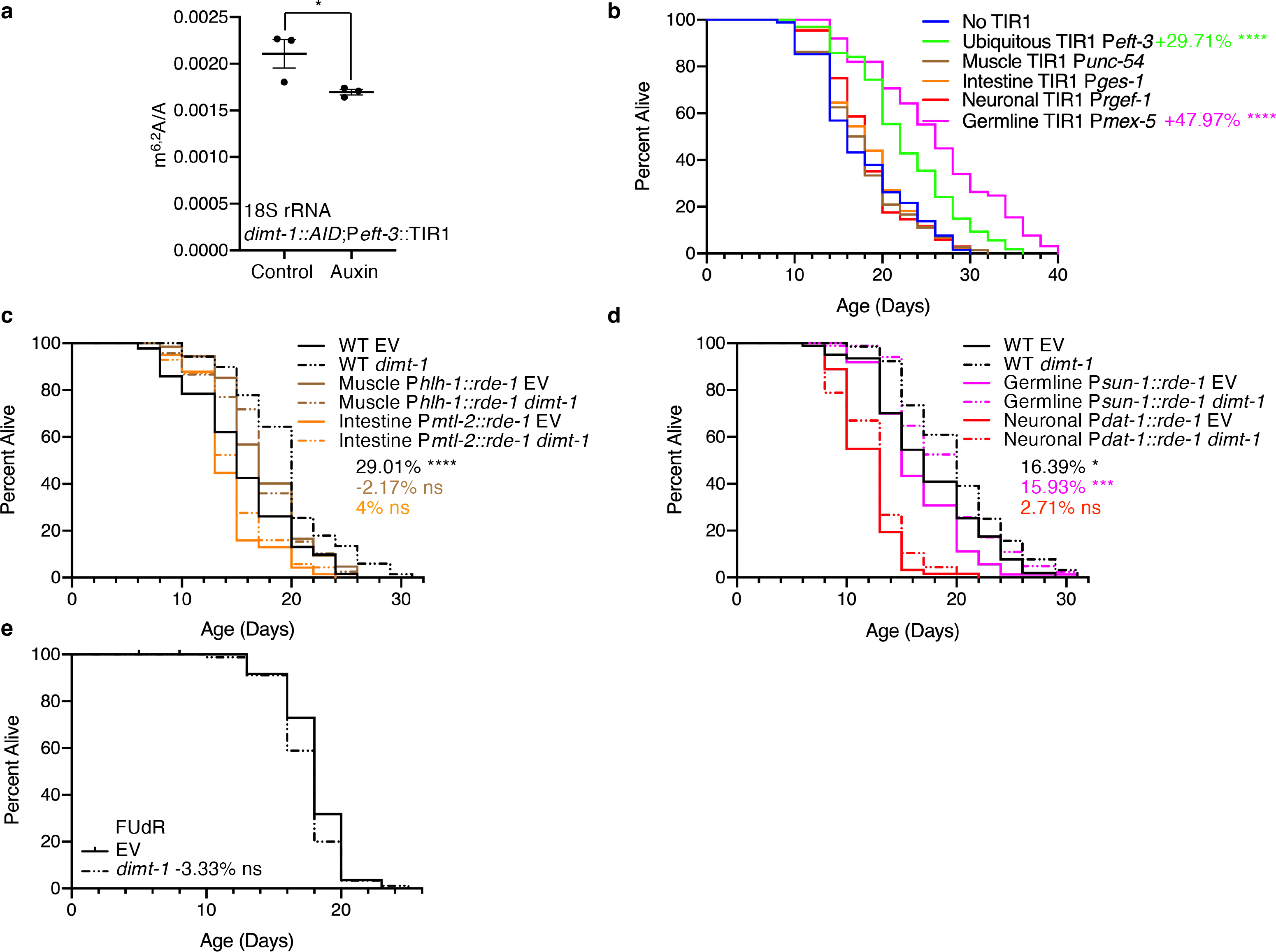
**DIMT-1 functions in the germline to regulate lifespan and affects translation of specific mRNA transcripts. a**, Auxin-inducible degron (AID) ubiquitous degradation of DIMT- 1 protein leads to significant decrease in m^6,2^A levels in the 18S rRNA subunit as assessed by UHPLC-ms/ms. **b,** Ubiquitous and germline-specific AID-induced DIMT-1 protein degradation causes a lifespan extension while DIMT-1 depletion in the muscle, intestine, or neurons has no effect on lifespan extension. **c-d,** Tissue-specific knock-down of *dimt-1* in the germline increases lifespan to a similar extent as in ubiquitous knock-down while knock-down of *dimt-1* in the muscle, intestine, or neurons has no significant effect on longevity. **e,** *dimt-1* knock-down does not extend the lifespan of worms treated with 5-fluorodeoxyuridine (FUdR), a drug that inhibits proliferation of germline stem cells and the production of intact eggs. Statistics and replicate longevity experiments are presented in Supplementary Table 2. ns; not-significant, *; p < 0.05, ***; p < 0.001, ****; p < 0.0001 as calculated by log-rank (mantel-cox) statistical test.

To validate our findings from the AID/TIR1 experiments, we performed an orthogonal approach by using tissue-specific RNAi strains to knock-down *dimt-1* and measure its effect on lifespan. When we specifically knocked down *dimt-1* in muscle, intestine, or neurons, we failed to observe a significant increase in lifespan; however, germline-specific knock-down of *dimt-1* caused a significant increase in lifespan that was comparable to knock-down of *dimt-1* in WT worms (16.93% in WT, 15.93% in germline-specific; p =0.8536 by 2-way ANOVA, **Fig. 4c, d**). Further bolstering the importance of the germline for the longevity effects of *dimt-1*, we found that treating worms with 5-fluorodeoxyuridine (FUdR), which inhibits proliferation of germline stem cells, the production of intact eggs in adults, and extends longevity^4,38^, abolished the effect of *dimt- 1* knock-down on lifespan (**Fig. 4e**).

## *dimt-1* affects ribosome binding to specific mRNA transcripts in the germline

We had previously found that knock-down of *dimt-1* in the parental generation caused a significant misregulation of both transcription and ribosome binding to transcripts involved in the determination of adult lifespan in the eggs of progeny^7^. To specifically examine which transcripts displayed altered binding by ribosomes after a decrease in DIMT-1 at later stages of life in the germline, we performed germline-specific Translating Ribosome Affinity Purification (TRAP)^39,40^ in four independent biological replicates from worms that were grown on empty vector (EV) control or *dimt-1* RNAi. We first validated that *dimt-1* knock-down extended longevity in this strain (Extended Data Fig. 3c). We next analyzed the transcription changes in response to *dimt-1* knock-down in post reproduction worms. Transcriptional changes would not be predicted to be direct consequences of manipulating the rRNA methylation, however, a natural consequence of changes in translation will lead to changes in transcription^41,42^. We found that 5,765 genes were differentially expressed, with 3,700 genes that were significantly upregulated and 2065 genes that were significantly downregulated upon *dimt-1* depletion (Extended Data Fig. 4a and Supplementary Table 3). Pathway analysis revealed increased expression of genes involved in longevity regulation, xenobiotic detoxifications, TGF-β, WNT, and MAPK signaling pathways, as well as degradation pathways including proteasome, peroxisome, and autophagy genes (Extended Data Fig. 4b, 4c, and Supplementary Table 3). Downregulated genes were enriched for protein processing, mTOR and FOXO signaling pathway, and ribosome biogenesis genes (Extended Data Fig. 4d, 4e, and Supplementary Table 3).

These altered transcriptional pathways could help to explain the extended longevity in response to *dimt-1* depletion, and the decreased expression of FOXO/DAF-16 and mTOR signaling pathways expression could help explain the *daf-16* and *raga-1* dependency of the lifespan extension (**Fig. 3f, g**). To determine if *dimt-1* depletion also altered the ribosome binding levels of specific transcripts, we sequenced ribosome-bound RNAs from the same four biological replicates isolated from the germline by TRAP and normalized that to the levels of the transcripts to measure translation efficiency. We found that 2082 genes were differently bound to ribosomes, with 1,666 genes that were significantly more bound and 416 genes that had significantly lower ribosome binding (**Fig. 5a** and Supplementary Table 4). Pathway analysis revealed that *dimt-1* depletion significantly altered the ribosomal binding to a selective subset of mRNAs involved in longevity regulation, degradation pathways, cellular detoxifications, glutathione metabolism, and oxidative phosphorylation (**Fig. 5b**, Extended Data Fig. 5, and Supplementary Table 4). The differences in ribosome bound transcripts could help to explain the altered longevity and stress resistance observed upon *dimt-1* depletion. Some of the pathways which were dysregulated on a transcriptional level were further dysregulated on a translational level as well while some other categories of genes appeared to only be misregulated transcriptionally or translationally as would be expected.

**Fig. 5.**
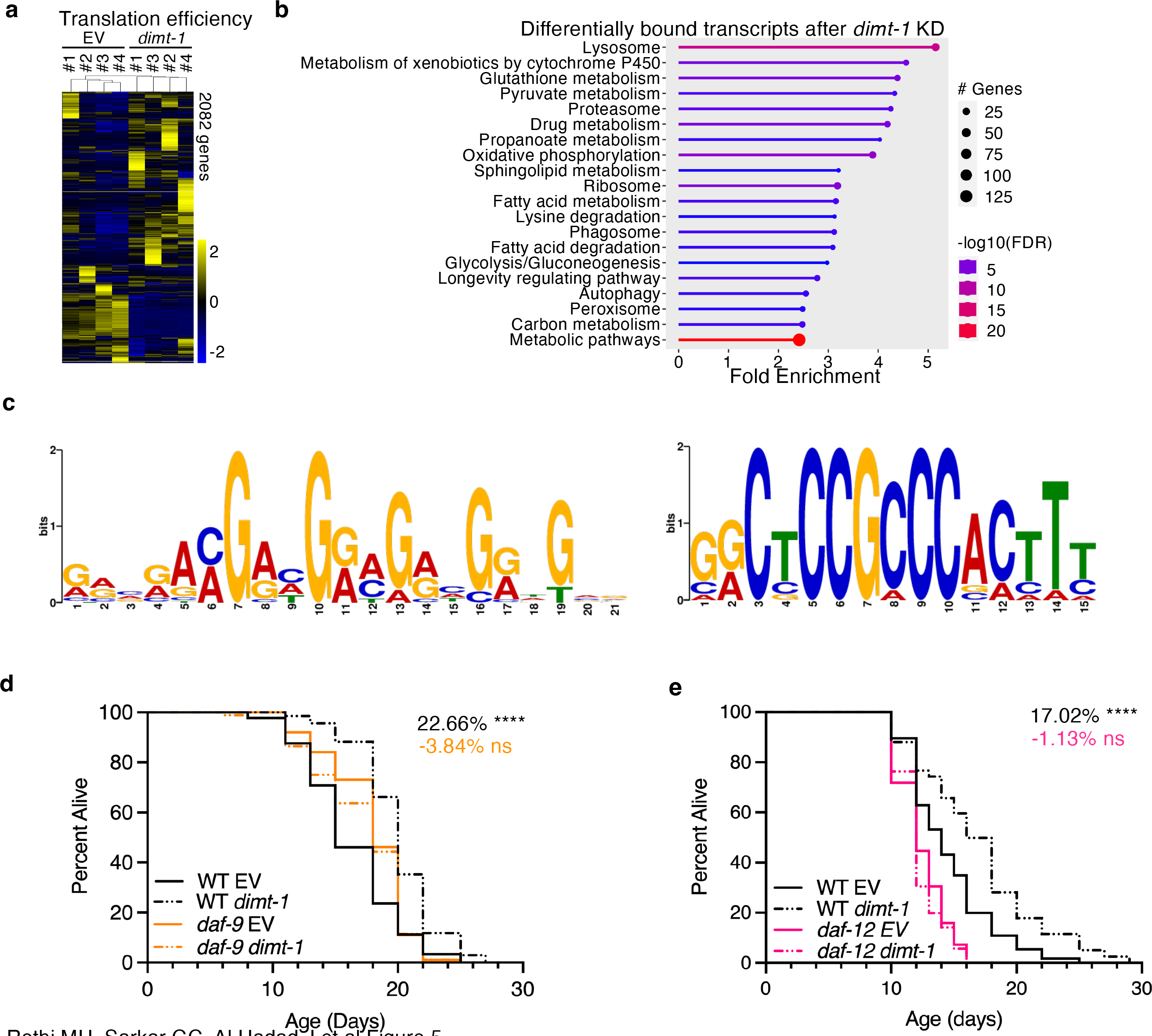
**DIMT-1 affects translation of specific mRNA transcripts. a**, Heat maps of the 2082 differentially ribosome bound transcripts in the ribosome after *dimt-1* knock-down from day 7 worms. Ribosome binding was normalized to total RNA expression to give translation efficiency. *Dimt-1* was knocked down from the L4 stage until day 7. Each column represents an independent biological replicate from ribosome sequencing after TRAP. **b,** Pathway analysis of differentially bound transcripts after *dimt-1* knock-down revealed altered ribosome binding to transcripts involved in longevity regulation, degradation, fatty acid metabolism, the ribosome, and oxidative phosphorylation. RNAseq and translation efficiency significantly regulated genes and gene ontology categories are presented in Supplementary Tables 3 and 4. **c,** Sequence motifs enriched in the 5’ UTR of more bound mRNA transcripts after *dimt-1* knock-down . **d,** *dimt-1* knock-down extends the lifespan of WT but not *daf-9* mutant worms (p=0.004 by 2-way ANOVA). **e,** *dimt-1* knock-down extends the lifespan of WT but not *daf-12* mutant worms (p=0.0005 by 2-way ANOVA).

Differentially binding to mRNA transcripts by ribosomes could be due to specific features present in the 5’UTR, such as specific sequences that may be translationally regulated^43,44^. To test if 18S m^6,2^A absence caused an enrichment of sequence motifs, we examined the 5’ UTR regions of mRNA transcripts which were ribosome bound when *dimt-1* was depleted in comparison to empty vector control. We identified 49 genes which have the sequence motif GRVRAMGAHGRMGRHGRWGVR and 11 genes which have the sequence motif GRCTCCGCCCACTTT (e-values 6.8E-4 and 5.4E4 respectively, **Fig. 5c**). This indicates that the presence of m^6,2^A on the 18S rRNA could regulate which transcripts are bound to the ribosome based partly on sequence features.

Interestingly one of the transcripts which displayed decreased ribosome occupancy when *dimt-1* was depleted was the cytochrome P450 enzyme, *daf-9.* DAF-9 has previously been demonstrated to produce a signaling lipid, dafachronic acid, which signals from the germline to the soma to activate the nuclear hormone receptor DAF-12 which inhibits organismal longevity^45–48^. We therefore tested whether the DAF-9/DAF-12 signaling pathway could be responsible for DIMT-1 regulated lifespan, as we found that DIMT-1 functions in the worm germline to regulate organismal lifespan. We found that *dimt-1* dependent lifespan extension was abolished in both *daf- 9* and *daf-12* mutant worms (**Fig. 5d, e**). This indicates that *daf-9* and *daf-12* are required for *dimt-1* induced lifespan extension. Taken together, these results suggest that *dimt-1* regulates organismal lifespan by selective translation in the germline of specific mRNA transcripts which subsequently leads to altered germline to soma signaling and lifespan.

## *dimt-1* regulates lifespan in later life

We were next interested in determining when DIMT-1 functioned to regulate organismal longevity. We performed a series of lifespan assays with AID-tagged *dimt-1* strains crossed with germline and ubiquitous TIR1 strains, where DIMT-1 was knocked out with auxin treatment at specific stages in the life of the worm (**Fig. 6a**). We found that when auxin was administered starting at the previous generation and for the entirety of the tested generation, starting at birth, or starting at the young adult stage, a consistent increase in lifespan was observed (52.19%, 41.33%, and 38.17% p<0.0001 in germline strain and 42.45%, 30.47%, and 35.45% p<0.0001 in ubiquitous strain, **Fig. 6b, c**). When auxin was only introduced from birth to the young adult stage and then worms were removed from auxin, we failed to observe a significant lifespan extension in the germline-specific DIMT-1 depletion strain (0.48% p=0.9570, **Fig. 5b**) and we only observed a modest lifespan extension in the ubiquitous DIMT-1 depletion strain (10.15% p=0.0101, **Fig. 6c**). Together, these results suggest that DIMT-1 functions in the germline after development to regulate lifespan. Although we determined that *dimt-1* regulates longevity following development, it was still unclear if the egg-laying phase is important for this phenotype, as we had previously found that DIMT-1 depletion causes a reduction in fertility^7^ and there is a well-known anti-correlation between reproduction and longevity^34^. We therefore examined the lifespan of our germline-specific TIR1 DIMT-1-AID tagged strain and introduced auxin at young adult, post egg-laying and mid-life phases. Surprisingly, we observed significant lifespan extension in all three stages compared to the untreated control (21.61%, 25.01%, and 18.93% respectively p<0.0001, **Fig. 6d**). This suggests that DIMT-1 affects lifespan post-developmentally, after egg-laying is complete, and in mid-life.

**Fig. 6.**
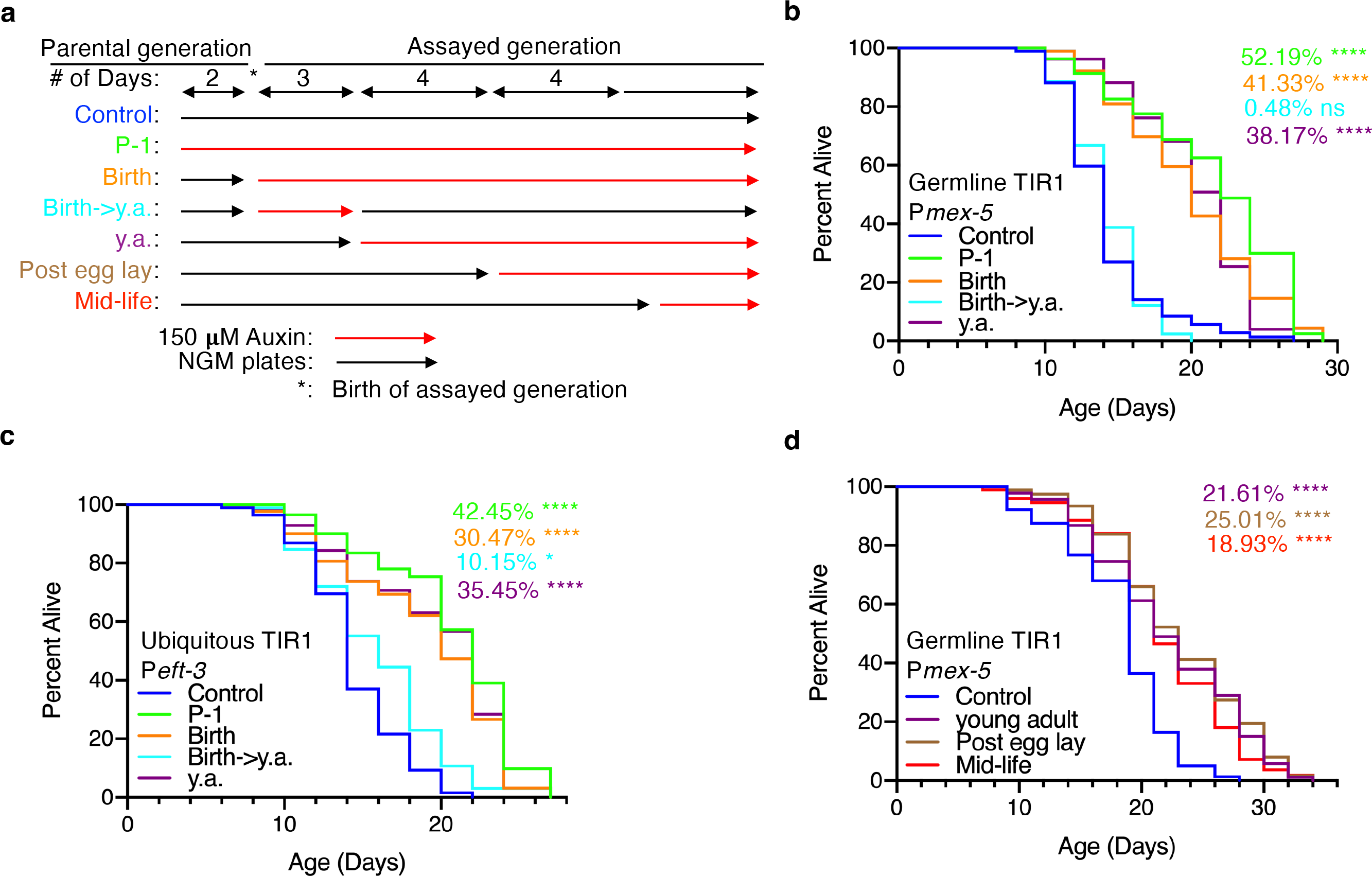
DIMT-1 regulates lifespan after mid-life. a, Experimental design for the AID-tagged DIMT-1 temporal knock out experiments (y.a. = young adults). Red lines indicate when strains were placed on 150 uM auxin. **b**, AID-induced depletion of DIMT-1 in the germline extends lifespan when depleted in the previous generation and for the entirety of the assayed generation, starting at birth, or from young adulthood for the remainder of the lifespan but does not extend lifespan when depleted only from birth until young adulthood. **c**, AID-induced depletion of DIMT- 1 ubiquitously extends lifespan when depleted in the previous generation and for the entirety of the assayed generation, starting at birth, or from young adulthood for the remainder of the lifespan and causes a less dramatic extension in lifespan when depleted only from birth until young adulthood. **d**, AID-induced depletion of DIMT-1 in the germline extends lifespan to a similar extent when depleted from young adulthood, after reproduction, or starting in mid-life. Statistics and replicate longevity experiments are presented in Supplementary Table 5. ns; not-significant, *; p < 0.05, ***; p < 0.001, ****; p < 0.0001 as calculated by log-rank (mantel-cox) statistical test.

## Discussion

Here, we show that DIMT-1, an evolutionarily conserved 18S rRNA m^6,2^A methyltransferase^7,9,10,49^, regulates lifespan in *C. elegans.* Reduction of DIMT-1 causes a lifespan extension that requires the DAF-16/FOXO transcription factor and TOR signaling. We found that DIMT-1 functions in the germline to regulate *C. elegans* lifespan and also causes differential binding of the ribosome to specific subsets of mRNA transcripts. One of the altered transcripts we identified was *daf-9* and we found that reduction of DIMT-1 caused a lifespan extension that required both the cytochrome P450 enzyme DAF-9 and its downstream nuclear hormone receptor transcription factor DAF-12. These results suggest an overall model whereby DIMT-1 regulates organismal lifespan by selective translation in the germline of specific mRNA transcripts which subsequently leads to altered germline to soma signaling and lifespan. Furthermore, while most longevity regulators identified to date function early in life to “lock-in” aging rates, we found that DIMT-1 can regulate lifespan after middle age. Together, this study provides evidence of selective translation, via ribosome heterogeneity, playing a significant role in the regulation of aging.

A major feature of aging is the loss of proteostasis, where maintenance of the appropriate amounts of proteins in the cell, is significantly disrupted later in life^50,51^. Previous studies have examined the levels of misfolded proteins which become increasingly prevalent in older organisms, causing breakdown of normal cellular function^52,53^. However, little is known about how transcripts are selected for translation during the aging process. Ribosomes were initially believed to be non-discriminatory, translating any mRNA transcripts they were presented with^54^, but recent work has suggested that specialized ribosomes can translate unique sets of transcripts under specific stress conditions^11,55–58^. We found that perturbations in many of the enzymes that regulate rRNA modifications have substantial effects on overall health and duration of lifespan in *C. elegans* (**Fig. 1**).

We^7^, and others^59^, have recently shown that DIMT-1’s function to N6-dimethylate two adjacent adenines on the 18S rRNA can direct the ribosome to specific subsets of transcripts for translation. Our previous work suggests that while DIMT-1 binding to rRNAs during rRNA processing is important for appropriate processing of rRNAs, the decreased expression of *dimt-1* that we see in response to *dimt-1* knock-down is not sufficient to significantly alter rRNA processing^7^. We also demonstrated that rRNAs are the predominant, if not the only, substrate of DIMT-1^7^. Here we additionally demonstrate that a complete ablation of DIMT-1’s catalytic activity is not lethal and even leads to increased lifespan and stress resistance (**Fig. 3a, b** and Extended Data Fig. 1d). These findings suggests that DIMT-1 is regulating lifespan through altering ribosome specialization, rather than through changing available amounts of processed rRNAs. In this work we found that reducing 18S rRNA m^6,2^A caused changes in ribosome binding in a set of selective mRNAs that are mostly involve in longevity, metabolisms, cellular detoxifications, protein homeostasis, degradation and recycling pathways specifically in the germline after reproduction.

Regulation of aging has been shown to involve both germline and somatic tissues^34^. However, most findings related to the germline regulation of lifespan have been shown to extend lifespan due to defects in germline development or reducing the energy diverted towards producing the next generation of progeny^34,60^. We showed that DIMT-1 functions through the germline, however, its role in lifespan extension takes place after development and reproduction (**Fig. 6**). This indicates that DIMT-1 is amenable to therapies or manipulation once the developmental or reproductive stage of an organism has passed.

It is interesting to note that global m^6,2^A levels decrease in whole worms as *C. elegans* age (**Fig. 2a**). This finding runs counter to the observation that *dimt-1* levels increase as *C. elegans* age (Extended Data Fig. 2a, b), and that depletion of *dimt-1* causes an increase in lifespan. Our working hypothesis to explain this apparent paradox is that this could reflect that m^6,2^A increases in some tissues or in some specific cells while it decreases in other tissues as organisms age. One other potential explanation for this apparent paradox could be that m^6,2^A could be important in responding to immediate environmental stresses in *C. elegans,* and rather than the absolute levels of m^6,2^A at particular ages being important for regulating lifespan, this modifications capacity to rapidly change could deteriorate with age. This notion is supported by the fact that m^6,2^A increases in response to both UV stress and heat stress (**Fig. 2b, c**). It will be interesting in future experiments to examine whether m^6,2^A levels increase specifically in the germline with age or if the capacity of this rRNA modification to respond to stresses later in life is diminished.

Several recent studies have demonstrated that there are proteomic, epigenetic, and epitranscriptomic changes associated with aging, and that enzymes which regulate these processes can also regulate aging^4,12,61–65^. Most regulators of aging determine organismal lifespan at early developmental stages^66–68^, however, several exceptions, including dietary restriction^69^ and this study, can increase lifespan later in life. Pathways which may be manipulated later in life to extend lifespan offer exciting possibilities to potentially address aging-associated diseases with interventions. Thus, altering rRNA modifications late in life may represent a new approach for addressing aging-related disorders and increasing health span. Nevertheless, our results provide evidence of selective translation, via ribosome heterogeneity, playing a significant role in the regulation of aging.

## Methods

### Strains used

N2 Bristol strain was used as the WT background. Worms were grown on *dam^-^dcm^-^* on standard nematode growth medium (NGM) plates in all experiments except for RNAi and auxin-inducible degron experiments. TIR1 expressing strains CA1200, DV3801, DV3803, DV3805, JDW221, and JDW225, the tissue specific RNAi strains DCL569, IGL1839, NR350, XE1474, the Flag tagged RPL-4 strain for germline specific ribosome purifications EV484, *eat-2(ad1116), daf-16(mu86), daf-2(e1370), hsf-1(sy441), clk-1(e2519), raga-1(ok386), glp-1(e2141ts), pgl-1(bn101ts), daf- 9(rh50),* and *daf-12(rh61rh412)* were obtained from the Caenorhabditis Genetics Center which is supported by the NIH office of research infrastructure programs P40OD010440. The AID-tagged DIMT-1 and the *dimt-1* E79A strains were generated by SunyBiotech. TIR1 strains were crossed with the AID-tagged DIMT-1 strain to generate double homozygous strains. Each strain was assayed by PCR to confirm the genotypes.

### Single-worm genotyping

Single worms were placed in 5 µl of worm lysis buffer [50 mM KCl, 10 mM Tris (pH 8.3), 2.5 mM MgCl2, 0.45% NP-40, 0.45% Tween 20, and proteinase K (60 mg/ml)] and incubated at -80°C for 1 hour, 60°C for 1 hour, and 95°C for 15 minutes. PCRs were performed using the following primers: E02H1.1 SY373 (forward), 5’-CGTCGAGGATGAGCGAGAAA-3’, E02H1.1 SY373 (reverse), 5’-TGGCCATTCCATTTTCATTACA-3’; DV3801, DV3803, DV3805, JDW221, JDW225 primers as described in (*40*); CA1200 primers as described in (*39*). PCRs were performed according to manufacturer’s protocol (New England Biolabs, M0273) and PCRs were resolved on agarose gels.

### Targeted RNAi screen

Bacteria expressing double-stranded RNA of putative ribosomal RNA methyltransferases were obtained from Blackwell lab, with bacteria carrying the empty vector backbone as negative control. Bacteria were grown at 37°C and seeded on NGM plates containing ampicillin (100 µg/ml) and isopropylthiogalactoside (IPTG; 0.4 mM). For experiments involving FuDR, bacteria were seeded on NGM plates containing ampicillin (100 µg/ml), FuDR (100mg/ml) and isopropylthiogalactoside (IPTG; 0.4 mM).

### Tissue specific and temporal knock-out with auxin inducible degron-tagged DIMT-1

NGM plates containing streptomycin (300 µg/ml) and 1-Napthaleneacetic acid (NAA, auxin, 150µM), were seeded with OP50-1 bacteria. Worms were grown on the auxin plates at specific worm stages for specific lengths of time according to the experiment shown in the results.

### Ultra-High Performance Liquid Chromatography coupled with Mass Spectrometry (UHPLC-MS/MS)

Total RNA was extracted by addition of 1 ml TRIzol (Invitrogen) to 100 µl of packed worm pellet. Six freeze-thaw cycles were performed in liquid nitrogen. RNA extraction was performed according to the manufacturer’s protocol (Invitrogen, TRIzol). To isolate 26S, 18S and 5.8/5S rRNAs, total RNA was run on agarose electrophoresis gels to separate rRNAs. rRNA bands were excised from the gel and purified with Zymoclean Gel RNA Recovery Kit (Zymo). For digestion to nucleosides, 250 ng of RNA samples were digested at 37°C for 2 hours with Nucleoside Digestion Mix (New England Biolabs, M069S). Digested RNA samples were diluted to 100 µl with double-distilled water and filtered through 0.22 µm Millex Syringe Filters. 5 µl of filtered solution was injected for LC-MS/MS analysis and analyzed using the Agilent 1290 UHPLC system with C18 reversed-phase column (2.1 mm by 50 mm; 1.8 m) as in^7^. Mobile Phase A is composed of water with 0.1% (v/v) formic acid, and mobile phase B is composed of acetonitrile with 0.1% (v/v) formic acid. MS detection was performed using an Agilent 6470 triple quadrupole mass spectrometer in positive electrospray ionization mode, and data was quantified in dynamic multiple reaction monitoring mode by monitoring mass transition 268 → 136 for adenosine, 282 → 136 for Am, 282 → 150 for m^6^A, 296 → 164 for m^6^2A, 284 → 152 for guanine, 298 → 152 for Gm, 298 → 166 for m^7^G, 244.1 → 112 for cytosine, 258 → 112 for Cm, 245 → 113 for uracil, 259 → 113 for Um, 245 → 125 for PseudoU.

### Longevity assays

Worm lifespan assays were performed at 20°C and 25°C. For longevity assays involving RNAi, worm populations were synchronized by placing L1 worms on NGM RNAi plates. Resulting eggs from the P^-1^ generation were transferred to new RNAi plates and hatching day for the P^0^ generation was counted as day 1 for all lifespan measurements. For longevity assay involving the auxin-inducible degron system, worm populations were synchronized by placing eggs for the P^0^ generation, on NGM plates seeded with OP50-1 bacteria either with or without 150µM auxin. Worms were changed every other day to new plates to avoid confounding progeny and were scored as dead or alive. Dead worms were scored if they did not respond to repeated prods with a platinum pick. Worms were censored if they died from vulval bursting or crawled off the plate. Each life- span assay used 90 worms on 3 plates (30 worms/plate). Data was plotted with Kaplan-Meier survival curves and statistical significance was tested using log-rank (Mantel-Cox) tests. Life-span assays were repeated at least once and showed similar trends in relative life-span effects.

### UV stress assays

For survival assays involving RNAi, the P^0^ generation was prepared as described in the longevity assays. Hatched eggs were allowed to grow to L4 stage on 3 plates per condition with 30 worms per plate (90 worms per assay). Worms were exposed to 0.8 Joules (J), then grown at 20°C, assessed every 24 hours for survival, and scored as dead or alive as described in the longevity assays. For UHPLC-MS/MS analysis, L4 worms were synchronized on NGM plates seeded with OP50-1 bacteria and exposed to 0.8 J. Then, the worms were allowed to recover for either 1 or 2 hours at 20°C, before being processed as described for the UHPLC-MS/MS analysis.

### Heat stress

For survival assays involving RNAi, the P^0^ generation was prepared as described in the longevity assays. Hatched eggs were allowed to grow to L4 stage on 3 plates per condition with 30 worms per plate (90 worms per assay). Worms were grown at 37°C for 6-7 hours, then grown at 20°C, assessed every 24 hours for survival, and scored as dead or alive as described in the longevity assays. For UHPLC-MS/MS analysis, L4 worms were synchronized on NGM plates seeded with OP50-1 bacteria and were grown at 37°C for 6-7 hours. Then the worms were allowed to recover for either 1 or 2 hours at 20°C, before being processed as described for the UHPLC-MS/MS analysis.

### Genetic epistasis

Specific worm strains as described in the results section, were grown on NGM RNAi plates seeded with either bacteria expressing double-stranded RNA for *dimt-1* gene or carrying the empty vector backbone as the negative control.

### Transcriptomic analysis

Transcriptomic analysis for the gene expression profiles of putative rRNA methyltransferases was performed using a publicly available dataset^19^.

### Translating Ribosome Affinity Purification (TRAP)

Synchronized EV484 (*efIs155[Cbr-unc-119(+) + Pmex-5::rpl-4::FLAG::tbb-2 3′UTR] II*) worms (germline-specific FLAG-tagged ribosomal protein *rpl-4*) were grown on NGM RNAi plates seeded with overnight grown HT1115 bacteria containing empty vector (control RNAi) till young adult stage. Young adult animals were than collected in M9 buffer and divided into two groups. Half of the worms were put into the fresh empty vector control RNAi plate and other half into the *dimt-1* RNAi plate for its knock-down (HT1115 bacteria containing a vector expressing double stranded RNA for *dimt-1* gene). In the background, we have also checked the lifespan extension phenotype of *dimt-1* using EV484 as a control experiment. Worms were grown at 20°C till the end of their reproductive phase (7-day old worms). Each day the P^0^ generation (parental) worms were filtered through a 20-micron mesh to remove confounding progenies during the egg-laying phase. On day 7, worms were washed thrice with M9 containing 1 mM cycloheximide (Sigma) and once with Lysis Buffer I (10 mM 4-(2-hydroxyethyl)-1-piperazineethanesulfonic acid (HEPES) pH 7.4, 150 mM KCl, 5 mM MgCl2, 0.5 mM DTT, 1 mM cycloheximide, Mini cOmplete Protease Inhibitor). Then, flash-frozen worm pearls were made using liquid nitrogen and homogenized using ice cold glass Dounce tissue homogenizer (around 30-40 strokes). Lysis Buffer II (Lysis Buffer I containing 0.5% v/v NP-40 (Sigma), 0.4 U/μL RNasin (Promega), 10 mM ribonucleoside vanadyl complex (RVC by NEB), 33 mM 1,2-diheptanoylsn-glycero-3-phosphocholine (DHPC by Merck) and 1% w/v sodium deoxycholate (Sigma)) was added to the sample and incubated on ice for 30 minutes. Samples were centrifuged for 12000 x rcf for 15 minutes at 4°C and the supernatant was collected (around 2 ml). 200 μl clear supernatant was collected for total RNA extraction (for mRNA sequencing). Remaining samples were added to washed Protein G coated DynaBeads for pre-clearing, incubated for 1 hour at 4°C rotating. Then, samples were incubated with 5 μL anti-FLAG antibody(1mg/ml)-coupled beads (F1804, sigma) for 2 hours at 4°C rotating and beads were washed 4 times with Wash Buffer (10 mM HEPES pH 7.4, 350 mM KCl, 5 mM MgCl2, 1% v/v NP-40). For RNA elution, the beads were incubated in 350 μL of RLT buffer (Qiagen RNeasy Kit) including 1% v/v β-mercaptoethanol for 10 min at room temperature. The eluate was separated from the beads and RNA was purified using RNeasy Plus Mini Kit (QIAGEN) according to manufacturer’s protocol.

### Transcriptome and ribosome profiling sequencing and analysis

RNA concentration was measured with Qubit using the RNA HS Assay kit. Libraries were prepared with TruSeq Stranded mRNA LT Sample Prep Kit according to TruSeq Stranded mRNA Sample Preparation Guide, Part # 15031047 Rev. E. Libraries were quantified using the Bioanalyzer (Agilent, Santa Clara, CA) and sequenced with Illumina NovaSeq6000 S4 (2×150bp) (reads trimmed to 2×100bp) to get 20M read depth coverage per sample. The BCL (binary base calls) files were converted into FASTQ using the Illumina package bcl2fastq. Fastq files were mapped to the WBCel235 *C. elegans* genome, and gene counts were obtained with STAR v2.7. 2b^70^. All subsequent steps were performed in R using WormBase gene IDs. After filtering of genes with low numbers of reads, the raw total mRNA-seq counts were normalized using edgeR^71^. Translation efficiency was calculated by dividing the raw ribosomal mRNA-seq counts by the raw total mRNA-seq counts. Differential gene expression analysis was performed using edgeR^71^ and was corrected for multiple comparisons using the Benjamini-Hochberg method. Statistical significance was defined as having a False Discovery Rate (FDR, or adjusted p-value) < 0.05. GSEA was performed using the clusterProfiler package^72^, heatmaps were generated using pheatmap package, and Revigo plots were generated using R code obtained from the Revigo webserver^73^. Sequence motif analysis for 5’ UTR regions of identified mRNA transcripts was performed using the MEME suite (https://meme-suite.org/meme/tools/meme)^74^.

## Acknowledgments

We thank J. Ward for advice about TIR1 strains. We thank T.K. Blackwell lab for RNAi clones. We thank members of the Greer laboratory for discussions and feedback on the manuscript. This work was supported by NIH grants (DP2AG055947, R56AG076496, and R01AG084739) to E.L.G and NIH grant T32 EB016652 to W.M.

## Author contributions

E.L.G. conceived and planned the study. M.H.R. and E.L.G. wrote the paper. M.H.R. helped complete Fig. 1, generated Figs. 2, 3e, 4a, 4e, 5c, 6b, 6c, S1b, S1d, and performed replicate lifespan assays in Table S1. J.A.H. produced Fig. 1, G.C.S. produced, isolated, and performed IPs of aged worms for ribosome sequencing, generated Figs. 4g, S3b-e, S2c, and S4, and performed lifespan assays, W.M. performed TRAP assay and ribosome sequencing analysis, generated Figs. 4F and S3A, and was advised by V.N.G., A.K.Y. helped conceive the project and optimize the UHPLC-ms/ms protocol and performed replicate lifespan assays, N.P. generated aging gradients for Fig. 2a and performed lifespan assays, R.S. performed replicate lifespan assays, J.N. helped produce Fig. 3c, S.D. helped M.H.R. isolate rRNAs and perform UHPLC-ms/ms, E.L.G. helped complete Fig. 1, generated Figs. 3a-d, 3f, 3g, 4b-d, 5, S1a, S1c,and S2a-b. All authors discussed the results and commented on the manuscript.

## Data deposition

Raw sequencing data can be accessed through the GEO repository, accession number GSE237802. Reviewers can access the data with the secure token: mnofcyoyhdytpwb. Bioinformatics pipelines and supplementary code are available at https://github.com/wtm09002/Greer_Dimt1.

## Competing interests

Authors declare no competing interests.

Correspondence and requests for materials should be addressed to

**Extended Data** Fig. 1**. DIMT-1 knock-down affects endoplasmic reticulum unfolding protein response (UPR^ER^). a**, No difference in HSP-16.2 expression in response to *dimt-1* knock-down as assessed by GFP fluorescence 5 hours after 30 minutes of heat shock at 37°C in *Phsp-16.2::gfp* (zSi3000) (UPR^cytosol^) worms. **b,** No difference in HSP-6 expression in response to *dimt-1* knock- down as assessed by GFP fluorescence after 5 hours of ethidium bromide (25 μg/mL) treatment in *Phsp-6::gfp* (zcIs13) (UPR^mitochondria^) worms. **c,** Decrease in HSP-4 expression in response to *dimt- 1* knock-down as assessed by GFP fluorescence after 5 hours of tunicamycin (5 μg/mL) treatment in *Phsp-4::gfp*(*zcIs4) (*UPR^ER^) worms. All quantification was done using ImageJ software. Statistics represent unpaired t-test with Welch’s correction. ns not significant, ** p < 0.01, **d,** Knock-down of *dimt-1* or mutation of the catalytic domain (E79A) caused increased survival on the ER stress inducer tunicamycin.

**Extended Data** Fig. 2**. Ribosomal RNA methyltransferases and modifications are dynamically regulated throughout life and change in response to heat stress. a,** Gene expression profile of putative rRNA methyltransferases in *C. elegans* from day 4 to day 19 age gradient. This data was generated by an analysis of gene expression data from^19^. **b,** *dimt-1* transcript levels increased with age relative to actin control as assessed by quantitative RT PCR. **c,** UHPLC-ms/ms analysis of rRNA modification levels in 26S and 18S rRNA subunits in an age gradient. Experiments were performed with 4 biological replicates. **d,** Gene expression profile of putative rRNA methyltransferases in *C. elegans* exposed to heat stress compared to control. This data was generated by an analysis of gene expression data from**^7^**. **e,** UHPLC-ms/ms analysis of rRNA modification levels in 26S and 18S rRNA subunits in worms exposed to heat stress compared to control. Experiments were performed with 4 biological replicates. **f,** UHPLC-ms/ms analysis of rRNA modification levels in 26S and 18S rRNA subunits in worms exposed to UV stress compared to control. Experiments were performed with 4 biological replicates. ns not significant, * p < 0.05, ** p < 0.01, *** p < 0.001, **** p < 0.0001.

**Extended Data** Fig. 3**. Lifespan extension by DIMT-1 does not require *clk-1* and occurs in the germline. a,** *dimt-1* knock-down increases in WT and *clk-1(e2519)* mutant worm lifespan to a similar extent (p=0.0855 by 2-way ANOVA). **b,** AID-induced DIMT-1 depletion extends lifespan in two different ubiquitous driver strains and when depleted specifically in the germline. **c,** *dimt-1* knock-down increases lifespan of EV484 mutant worms. Statistics and replicate experiments are presented in Supplementary Tables 1 and 2. ns; not-significant, *; p < 0.05, **; p < 0.01, ***; p < 0.001, ****; p < 0.0001 as calculated by log-rank (mantel-cox) statistical test.

**Extended Data** Fig. 4**. *dimt-1* depletion causes a misregulation of expression of genes involved in longevity regulation, MTOR and FoxO/DAF-16 signaling and degradation pathways. a,** A heat map of 5,765 differentially transcribed genes in response to *dimt-1* knock-down in day 7 worms. *dimt-1* was knocked down from the L4 stage until day 7. Each column represents an independent biological replicate. **b-c,** Pathway analysis of genes which show increased expression in response to *dimt-1* knock-down reveal genes involved in longevity regulation, TGF-β, WNT and MAPK signaling pathways, as well as degradation pathways including proteasome, peroxisome, and autophagy genes. **d-e,** Pathway analysis of genes which showed decreased expression in response to *dimt-1* knock-down reveal genes involved in protein processing, MTOR and FoxO signaling pathway, and ribosome biogenesis genes. Gene details and gene ontology pathways is presented in Supplementary Table 3.

**Extended Data** Fig. 5**. *dimt-1* depletion causes altered binding of the ribosome to transcripts involved in longevity regulation, oxidative phosphorylation, degradation pathways, and glutathione metabolism in the germline of day 7 worms. a,** Pathway analysis of 2082 differentially bound transcripts in the germline of day 7 worms treated with *dimt-1* RNAi from the L4 stage until day 7 reveals transcripts involved in longevity regulation, oxidative phosphorylation, degradation pathways and glutathione metabolism. **b-c,** Pathway analysis of transcripts that show increased binding in response to *dimt-1* depletion include genes involved in longevity regulation, fatty acid metabolism, glutathione metabolism, metabolism, and various degradation pathways. **d-e,** Pathway analysis of transcripts that show decreased binding in response to *dimt-1* depletion in the germline include genes involved in the ribosome, proteasome, oxidative phosphorylation and longevity regulation. Gene details and gene ontology pathways is presented in Supplementary Table 4.

**Supplementary Table 1. Dimt-1 depletion extends lifespan in a raga-1 and germline dependent manner** The figure panels in which specific experiments are shown or used are indicated in the right column. The mean lifespan and SD values were calculated by Prism from triplicate samples of 30 worms each (90 worms total). # worms: number of observed dead worms at the end of the experiment/number of alive worms at the beginning of the experiment. The difference between both numbers corresponds to the number of censored worms (worms that underwent “matricide”, exhibited ruptured vulva, or crawled off the plates). P values are calculated by log rank (Mantel-Cox) statistical test.

**Supplementary Table 2. DIMT-1 functions in the germline to regulate lifespan** The figure panels in which specific experiments are shown or used are indicated in the right column. The mean lifespan and SD values were calculated by Prism from triplicate samples of 30 worms each (90 worms total). # worms: number of observed dead worms at the end of the experiment/number of alive worms at the beginning of the experiment. The difference between both numbers corresponds to the number of censored worms (worms that underwent “matricide”, exhibited ruptured vulva, or crawled off the plates). P values are calculated by log rank (Mantel-Cox) statistical test.

**Supplementary Table 5. DIMT-1 functions after mid-life to regulate lifespan** The figure panels in which specific experiments are shown or used are indicated in the right column. The mean lifespan and SD values were calculated by Prism from triplicate samples of 30 worms each (90 worms total). # worms: number of observed dead worms at the end of the experiment/number of alive worms at the beginning of the experiment. The difference between both numbers corresponds to the number of censored worms (worms that underwent “matricide”, exhibited ruptured vulva, or crawled off the plates). P values are calculated by log rank (Mantel-Cox) statistical test.

