## Supplemental Figures 1-5 Supplemental Tables 1,2,+5 for "The 18S rRNA Methyltransferase DIMT-1 Regulates Lifespan in the Germline Later in Life"

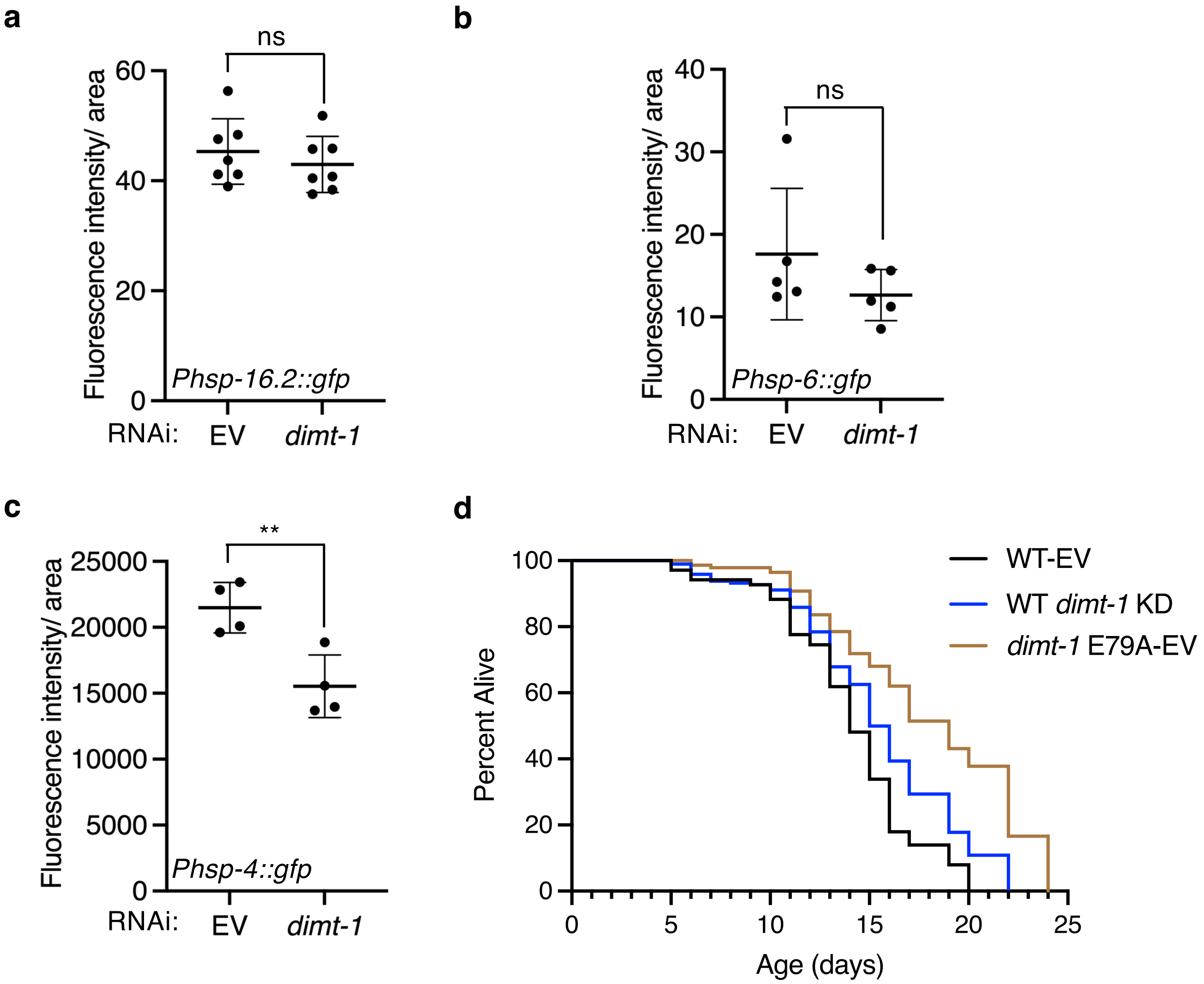


Rothi MH, Sarkar GC, Al Hadad J et al Fig S1

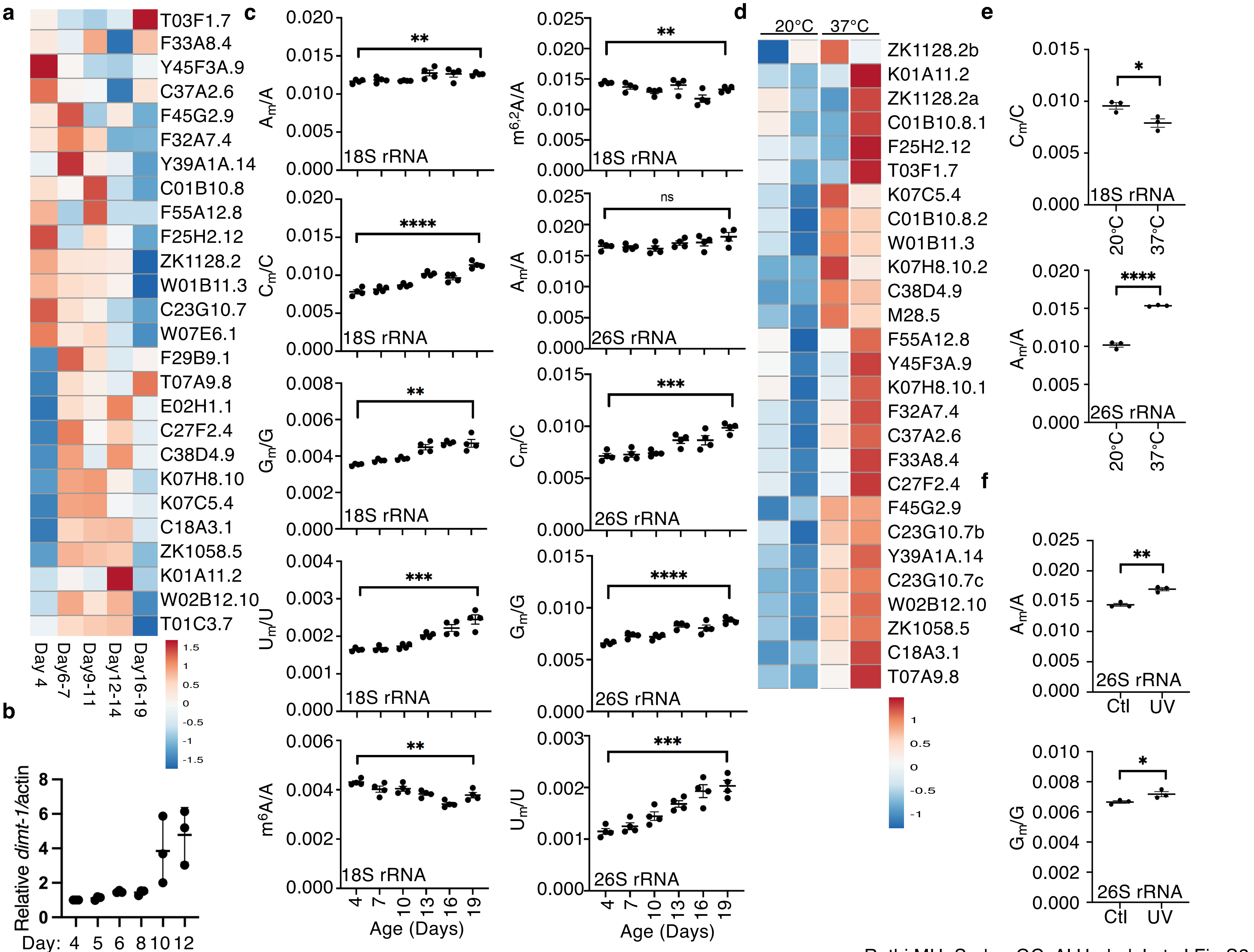


Rothi MH, Sarkar GC, Al Hadad J et al Fig S2

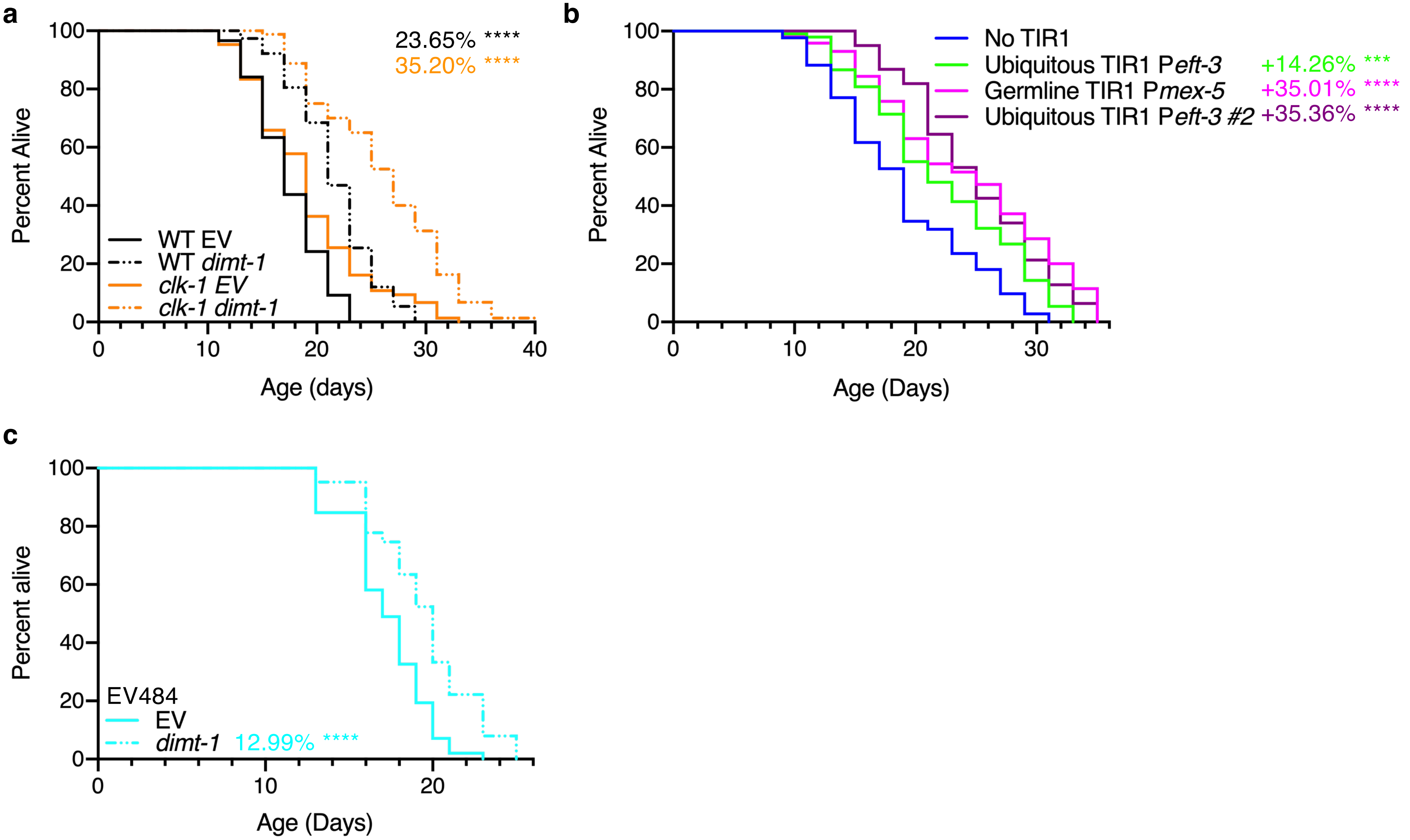


Rothi MH, Sarkar GC, Al Hadad J et al Fig S3

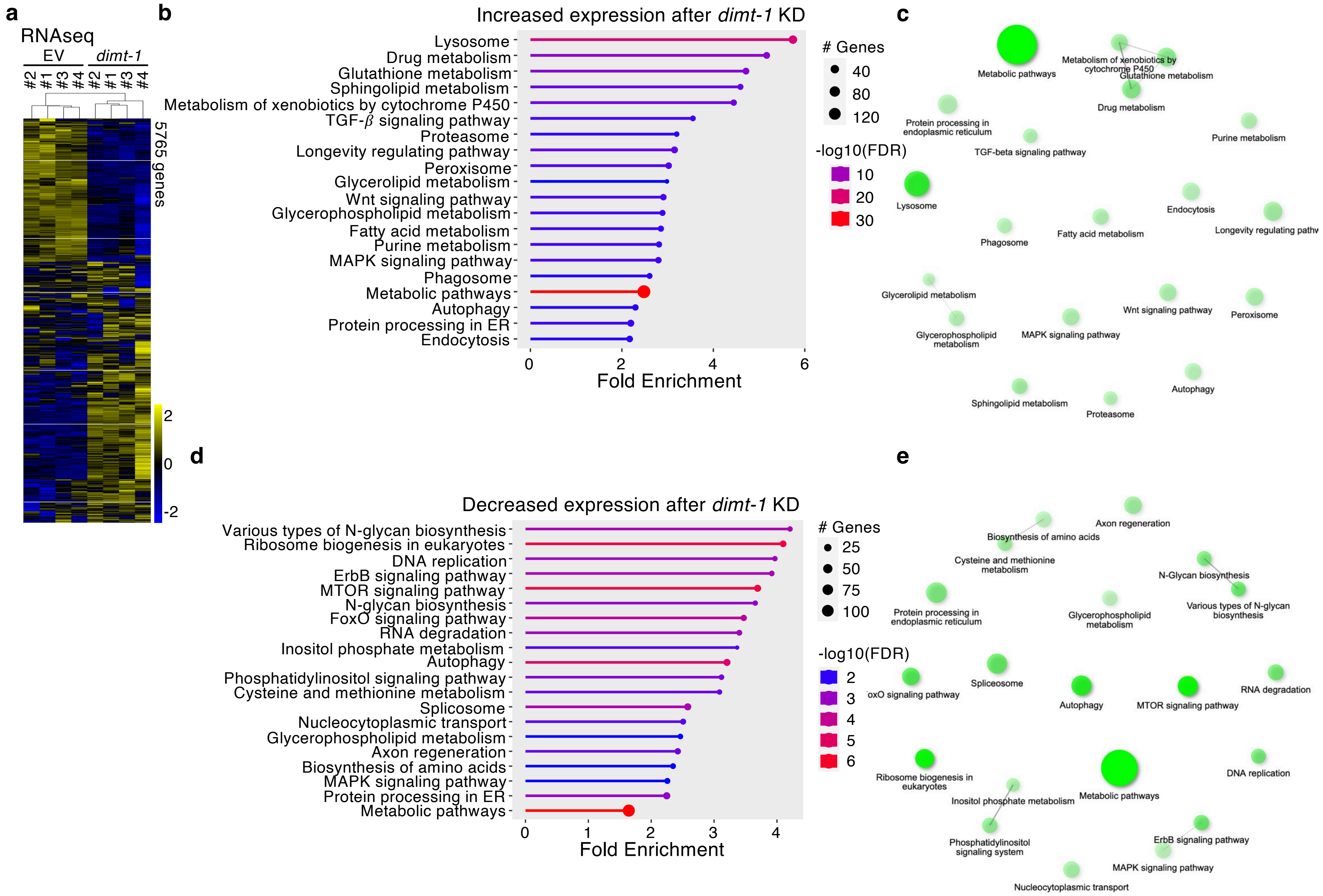


Rothi MH, Sarkar GC, Al Hadad J et al Fig S4

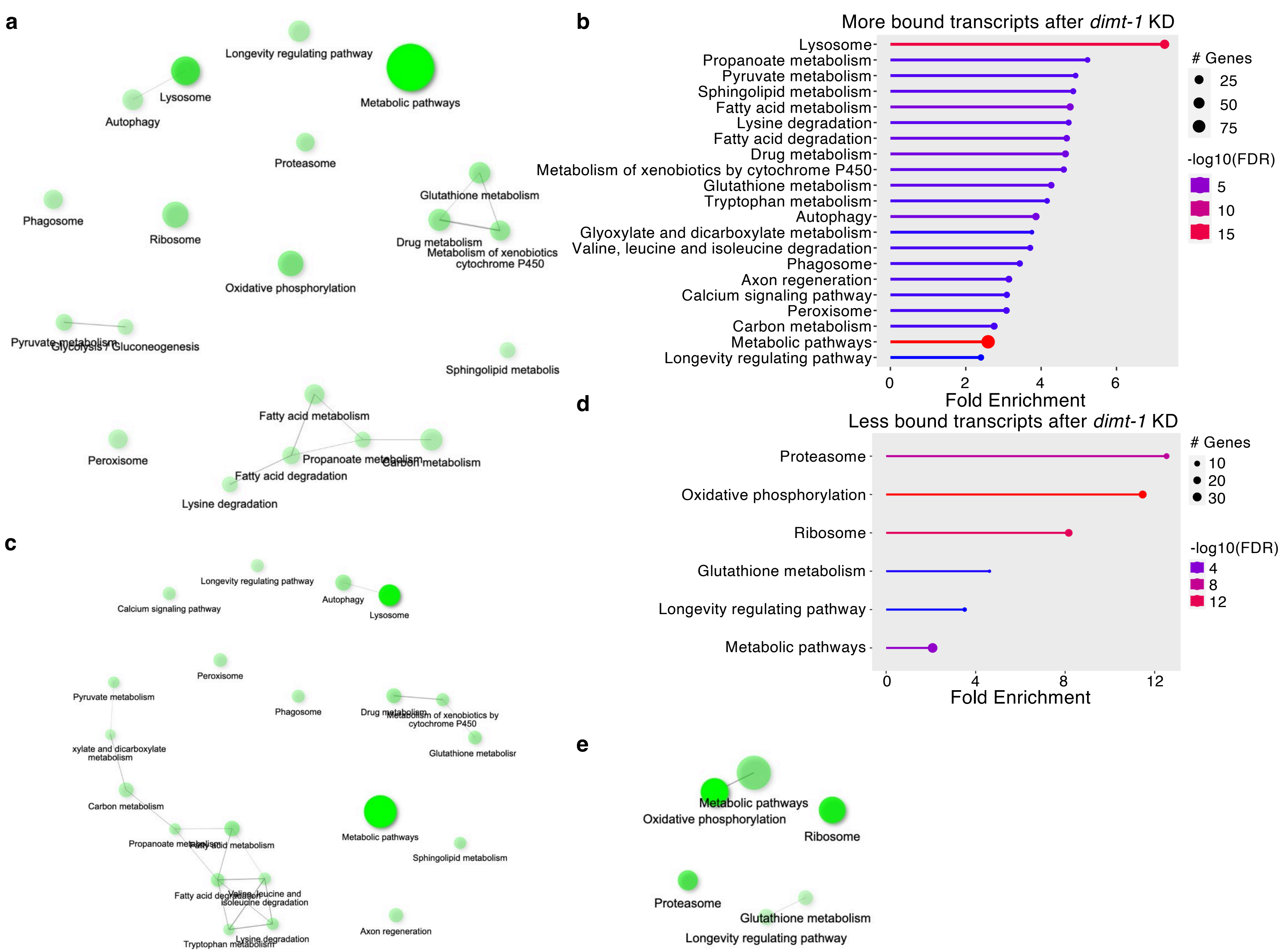


Rothi MH, Sarkar GC, Al Hadad J et al Fig S5

| Strain | RNAi | Mean +/- SEM | Median | p values | # worms | Figure |
|---|---|---|---|---|---|---|
| WT | EV | 16.211 +/- 0.3620 | 17 | | 128/163 | 3E |
| WT | *dimt-1* | 17.949 +/- 0.4724 | 17 | 0.0012 | 117/172 | 3E |
| *hsf-1* | EV | 14.714 +/- 0.5969 | 15 | | 49/70 | 3E |
| *hsf-1* | *dimt-1* | 17.951 +/- 0.5677 | 17 | 0.0031 | 68/87 | 3E |
| *daf-16(mu86)* | EV | 15.279 +/- 0.4190 | 15 | | 68/82 | |
| *daf-16(mu86)* | *dimt-1* | 15.921 +/- 0.2927 | 17 | 0.3626 | 63/90 | |
| *eat-2(ad1116)* | EV | 23.433 +/- 1.318 | 24 | | 60/88 | |
| *eat-2(ad1116)* | *dimt-1* | 26.286 +/- 1.249 | 28 | 0.6196 | 35/91 | |
| *glp-1(e2141ts)* | EV | 21.258 +/- 0.6080 | 21 | | 89/91 | |
| *glp-1(e2141ts)* | *dimt-1* | 22.943 +/- 0.7214 | 24 | 0.0189 | 88/95 | |
| WT | EV | 17.967 +/- 0.3376 | 19 | | 90/92 | |
| WT | *dimt-1* | 20.208 +/- 0.4085 | 21 | <0.0001 | 77/88 | |
| *daf-16(mu86)* | EV | 16.5 +/- 0.3545 | 17 | | 88/88 | |
| *daf-16(mu86)* | *dimt-1* | 18.053 +/- 0.3685 | 19 | 0.0017 | 75/79 | |
| *eat-2(ad1116)* | EV | 23.728 +/- 0.6403 | 21 | | 92/96 | |
| *eat-2(ad1116)* | *dimt-1* | 25.639 +/- 0.6912 | 26 | 0.0574 | 83/87 | |
| *raga-1* | EV | 27.242 +/- 0.7933 | 28 | | 95/95 | |
| *raga-1* | *dimt-1* | 26.659 +/- 0.7021 | 26 | 0.5407 | 86/87 | |
| *daf-2(e1370)* | EV | 38.029 +/- 1.435 | 41 | | 70/97 | |
| *daf-2(e1370)* | *dimt-1* | 48.75 +/- 2.028 | 51 | <0.0001 | 61/94 | |
| *glp-1(e2141ts)* | EV | 26.69 +/- 0.7349 | 28 | | 100/102 | |
| *glp-1(e2141ts)* | *dimt-1* | 22.553 +/- 0.5352 | 23 | <0.0001 | 103/104 | |
| WT | EV | 17.414 +/- 0.3643 | 17 | | 87/96 | 3C,3F,S3A |
| WT | *dimt-1* | 21.533 +/- 0.4659 | 21 | <0.0001 | 75/94 | 3C,3F,S3A |
| *daf-16(mu86)* | EV | 15.230 +/- 0.3720 | 15 | 0.0001 | 87/96 | 3F |
| *daf-16(mu86)* | *dimt-1* | 16.012 +/- 0.4390 | 15 | 0.0907 | 81/96 | 3F |
| *eat-2(ad1116)* | EV | 20.986 +/- 0.7766 | 7 | <0.0001 | 72/96 | 3C |
| *eat-2(ad1116)* | *dimt-1* | 27.397 +/- 0.5261 | 19 | <0.0001 | 68/74 | 3C |
| *hsf-1* | EV | 7.8 +/- 0.6208 | 19 | <0.0001 | 45/76 | |
| *hsf-1* | *dimt-1* | 17.375 +/- 0.6098 | 27 | <0.0001 | 59/66 | |
| *clk-1* | EV | 19.107 +/- 0.6696 | 19 | 0.0042 | 75/96 | S3A |
| *clk-1* | *dimt-1* | 25.833 +/- 0.7145 | 27 | <0.0001 | 78/96 | S3A |
| WT | EV | 15.181 +/- 0.4120 | 14 | | 94/98 | 3H |
| WT | *dimt-1* | 19.481 +/- 0.5768 | 20 | <0.0001 | 81/91 | 3H |
| *glp-1(e2141ts)* | EV | 26.259 +/- 0.6842 | 26 | <0.0001 | 81/82 | 3H |
| *glp-1(e2141ts)* | *dimt-1* | 24.885 +/- 0.7855 | 24 | 0.2013 | 61/73 | 3H |
| *pgl-1* | EV | 18.443 +/- 0.9143 | 16 | <0.0001 | 97/98 | |
| *pgl-1* | *dimt-1* | 17.674 +/- 0.7517 | 18 | 0.2133 | 92/97 | |
| WT | EV | 17.90 +/- 0.2304 | 18 | | 83/96 | |
| WT | *dimt-1* | 21.64 +/- 0.4192 | 23 | <0.0001 | 81/96 | |
| WT | EV | 19.45 +/- 0.4091 | 21 | | 65/91 | |
| WT | *dimt-1* | 26.86 +/- 1.269 | 24 | 0.0003 | 79/89 | |
| *raga-1* | EV | 27.60 +/- 0.9856 | 31 | <0.0001 | 43/88 | |
| *raga-1* | *dimt-1* | 30.03 +/- 1.260 | 27 | 0.4212 | 65/89 | |

**Supplementary Table 1. Dimt-1 depletion extends lifespan in a raga-1 and germline dependent manner** The figure panels in which specific experiments are shown or used are indicated in the right column. The mean lifespan and SD values were calculated by Prism from triplicate samples of 30 worms each (90 worms total). # worms: number of observed dead worms at the end of the experiment/number of alive worms at the beginning of the experiment. The difference between both numbers corresponds to the number of censored worms (worms that underwent “matricide”, exhibited ruptured vulva, or crawled off the plates). P values are calculated by log rank (Mantel-Cox) statistical test.

| Strain | RNAi | Mean +/- SD | Median | p values | # worms | Figure |
|---|---|---|---|---|---|---|
| WT | EV | 17.778 +/- 0.4064 | 17 | | 90/94 | 3D, 3I |
| WT | *dimt-1* | 24.918 +/- 0.6084 | 27 | <0.0001 | 50/60 | 3D, 3I |
| *pgl-1* | EV | 18.449 +/- 0.4319 | 20 | 0.2051 | 99/99 | 3I |
| *pgl-1* | *dimt-1* | 19.065 +/- 0.4880 | 20 | 0.2807 | 94/94 | 3I |
| *daf-2(e1370)* | EV | 38.548 +/- 1.974 | 41 | <0.0001 | 31/45 | 3D |
| *daf-2(e1370)* | *dimt-1* | 51.647 +/- 2.980 | 55 | <0.0001 | 34/68 | 3D |
| WT FUdR | EV | 17.581 +/- 0.3725 | 19 | | 83/92 | |
| WT FUdR | *dimt-1* | 15.709 +/- 0.3695 | 17 | <0.0001 | 87/94 | |
| WT FUdR | EV | 17.953 +/- 0.2396 | 18 | | 85/92 | 4E |
| WT FUdR | *dimt-1* | 17.400 +/- 0.2452 | 18 | 0.0773 | 90/93 | 4E |
| WT | EV | 16.281 +/- 0.5470 | 15 | | 90/92 | 3G, 5D |
| WT | *dimt-1* | 19.971 +/- 0.9724 | 20 | <0.0001 | 77/88 | 3G, 5D |
| *daf-16(mu86)* | EV | 12.943 +/- 0.7447 | 13 | | 88/88 | |
| *daf-16(mu86)* | *dimt-1* | 12.964 +/- 0.7372 | 13 | 0.9641 | 75/79 | |
| *daf-9(rh50)* | EV | 17.88 +/- 0.7043 | 18 | | 92/96 | 5D |
| *daf-9(rh50)* | *dimt-1* | 17.193 +/- 0.7995 | 18 | 0.5511 | 83/87 | 5D |
| *raga-1* | EV | 24.402 +/- 0.7772 | 22 | | 95/95 | 3G |
| *raga-1* | *dimt-1* | 24.531 +/- 0.7260 | 22 | 0.7892 | 86/87 | 3G |
| WT | EV | 14.51 +/- 0.42 | | | 67/90 | 5E |
| WT | *dimt-1* | 16.98 +/- 0.52 | | 0.0002 | 94/94 | 5E |
| *daf-12(rh61rh412)* | EV | 12.42 +/- 0.20 | | | 96/96 | 5E |
| *daf-12(rh61rh412)* | *dimt-1* | 12.28 +/- 0.20 | | 0.57 | 81/90 | 5E |
| *daf-9(rh50)* | EV | 13.22 +/- 0.19 | | | 94/90 | |
| *daf-9(rh50)* | *dimt-1* | 12.84 +/- 0.22 | | 0.34 | 80/90 | |
| WT | EV | 16.43 +/- 0.58 | | | 76/90 | |
| WT | *dimt-1* | 19.31 +/- 0.45 | | <0.0001 | 140/140 | |
| *daf-12(rh61rh412)* | EV | 13.74 +/- 0.46 | | | 79/90 | |
| *daf-12(rh61rh412)* | *dimt-1* | 13.86 +/- 0.47 | | 0.89 | 74/90 | |
| *daf-9(rh50)* | EV | 13.7 +/- 0.46 | | | 65/90 | |
| *daf-9(rh50)* | *dimt-1* | 13.53 +/- 0.38 | | 0.87 | 102/102 | |
| WT | | 15.512 +/- 1.992 | 15 | | 83/96 | 3B |
| *dimt-1* E79A | | 21.637 +/- 0.2514 | 22 | <0.0001 | 81/96 | 3B |
| WT | | 15.383 +/- 1.899 | 15 | | 83/96 | |
| *dimt-1* E79A | | 21.3 +/- 0.2638 | 20 | <0.0001 | 81/96 | |
| WT | | 15.598 +/- 1.703 | 15 | | 83/96 | |
| *dimt-1* E79A | | 20.883 +/- 0.3052 | 20 | <0.0001 | 81/96 | |

**Supplementary Table 1. Dimt-1 depletion extends lifespan in a raga-1 and germline dependent manner** The figure panels in which specific experiments are shown or used are indicated in the right column. The mean lifespan and SD values were calculated by Prism from triplicate samples of 30 worms each (90 worms total). # worms: number of observed dead worms at the end of the experiment/number of alive worms at the beginning of the experiment. The difference between both numbers corresponds to the number of censored worms (worms that underwent “matricide”, exhibited ruptured vulva, or crawled off the plates). P values are calculated by log rank (Mantel-Cox) statistical test.

| Strain | TIR1 | Time Placed on auxin | Mean +/- SD | Median | p values | # worms | Figure |
|---|---|---|---|---|---|---|---|
| WT | | P-1 L4 | 16.233 +/- 0.4988 | 16 | 0.0746 | 60/93 | |
| *dimt-1::AID* | | P-1 L4 | 17.123 +/- 0.6470 | 16 | | 73/96 | 4A |
| *dimt-1::AID* | CA1200 eft-3 ubiquitous | P-1 L4 | 22.211 +/- 0.8067 | 22 | <0.0001 | 57/85 | 4A |
| *dimt-1::AID* | DV3801 unc-54 muscle | P-1 L4 | 17.36 +/- 0.6112 | 17 | 0.9993 | 75/92 | 4A |
| *dimt-1::AID* | DV3803 ges-1 intestine | P-1 L4 | 17.922 +/- 0.6650 | 18 | 0.5125 | 51/88 | 4A |
| *dimt-1::AID* | DV3805 rgef-1 neuron | P-1 L4 | 17.85 +/- 0.6920 | 18 | 0.6655 | 40/75 | 4A |
| *dimt-1::AID* | JDW221 mex-5 germline | P-1 L4 | 25.351 +/- 0.8675 | 26 | <0.0001 | 74/97 | 4A |
| WT | | P-1 L4 | 16.86 +/- 0.4803 | 17 | | 71/90 | |
| *dimt-1::AID* | | P-1 L4 | 18.66 +/- 0.6840 | 19 | | 76/90 | S2B |
| *dimt-1::AID* | CA1200 eft-3 ubiquitous | P-1 L4 | 21.03 +/- 0.7266 | 21 | 0.0007 | 74/103 | S2B |
| *dimt-1::AID* | DV3801 unc-54 muscle | P-1 L4 | 17.31 +/- 0.6508 | 17 | 0.2456 | 72/94 | |
| *dimt-1::AID* | DV3803 ges-1 intestine | P-1 L4 | 17.17 +/- 0.7407 | 17 | 0.2780 | 69/91 | |
| *dimt-1::AID* | DV3805 rgef-1 neuron | P-1 L4 | 17.34 +/- 0.6514 | 15 | 0.1238 | 77/95 | |
| *dimt-1::AID* | JDW221 mex-5 germline | P-1 L4 | 24.20 +/- 0.8902 | 25 | <0.0001 | 70/73 | S2B |
| *dimt-1::AID* | JDW225 eft-3 ubiquitous | egg | 24.42 +/- 0.7870 | 25 | <0.0001 | 52/18 | S2B |

| Strain | RNAi | Mean +/- SD | Median | p values | # worms | Figure |
|---|---|---|---|---|---|---|
| WT | EV | 15.063 +/- 0.6362 | 15 | | 64/90 | 4C |
| WT | *dimt-1* | 19.433 +/- 0.5664 | 20 | <0.0001 | 67/90 | 4C |
| NR350 | EV | 17.326 +/- 0.6276 | 17 | | 43/90 | 4C |
| NR350 | *dimt-1* | 16.95 +/- 0.7071 | 17 | 0.8636 | 40/90 | 4C |
| DCL569 | EV | 16.222 +/- 0.5706 | 15 | | 72/90 | |
| DCL569 | *dimt-1* | 17.649 +/- 0.5898 | 17 | 0.0924 | 74/90 | |
| IG1836 | EV | 14.2 +/- 0.3896 | 13 | | 70/98 | 4C |
| IG1836 | *dimt-1* | 14.768 +/- 0.4562 | 15 | 0.2763 | 69/90 | 4C |
| XE14781 | EV | 12.086 +/- 0.4714 | 10 | | 58/90 | |
| XE14781 | *dimt-1* | 12.413 +/- 0.3590 | 13 | 0.6165 | 63/90 | |
| WT | EV | 17.170 +/- 0.7227 | 17 | | 53/90 | 4D |
| WT | *dimt-1* | 19.984 +/- 0.6028 | 20 | 0.0134 | 64/90 | 4D |
| NR350 | EV | 16.923 +/- 0.7433 | 17 | | 39/90 | |
| NR350 | *dimt-1* | 18.412 +/- 0.6861 | 18.5 | 0.2357 | 34/90 | |
| DCL569 | EV | 16.347 +/-0.4772 | 15 | | 72/90 | 4D |
| DCL569 | *dimt-1* | 18.951 +/- 0.4957 | 20 | 0.0004 | 82/90 | 4D |
| IG1836 | EV | 15.226 +/- 0.4148 | 15 | | 62/90 | |
| IG1836 | *dimt-1* | 14.761 +/- 0.4563 | 13 | 0.4794 | 71/90 | |
| XE14781 | EV | 11.952 +/- 0.3383 | 13 | | 62/90 | 4D |
| XE14781 | *dimt-1* | 12.412 +/- 0.3859 | 13 | 0.2677 | 68/90 | 4D |
| EV484 | EV | 18.30 +/- 0.25 | 18 | | 47/53 | |
| EV484 | *dimt-1* | 19.10 +/- 0.46 | 20 | 0.0141 | 58/67 | |
| EV484 | EV | 18.38 +/- 0.27 | 19 | | 76/85 | |
| EV484 | *dimt-1* | 17.58 +/- 0.37 | 18 | 0.1787 | 54/60 | |
| EV484 | EV | 16.49 +/- 0.30 | 17 | | 49/54 | |
| EV484 | *dimt-1* | 18.82 +/- 0.33 | 19 | <0.0001 | 49/53 | |
| EV484 | EV | 17.85 +/- 0.24 | 18 | | 62/67 | |
| EV484 | *dimt-1* | 19.75 +/- 0.40 | 20 | <0.0001 | 40/46 | |
| EV484 | EV | 17.24 +/- 0.25 | 17 | | 98/107 | S2C |
| EV484 | *dimt-1* | 19.48 +/- 0.38 | 20 | <0.0001 | 63/65 | S2C |

**Supplementary Table 2. DIMT-1 functions in the germline to regulate lifespan** The figure panels in which specific experiments are shown or used are indicated in the right column. The mean lifespan and SD values were calculated by Prism from triplicate samples of 30 worms each (90 worms total). # worms: number of observed dead worms at the end of the experiment/number of alive worms at the beginning of the experiment. The difference between both numbers corresponds to the number of censored worms (worms that underwent "matricide", exhibited ruptured vulva, or crawled off the plates). P values are calculated by log rank (Mantel-Cox) statistical test.

| Strain | TIR1 | Time Placed on auxin | Mean +/- SD | Median | p values | # worms | Figure |
|---|---|---|---|---|---|---|---|
| *dimt-1::AID* | JDW221 mex-5 germline | | 14.127 +/- 0.4027 | 14 | | 72/90 | |
| *dimt-1::AID* | JDW221 mex-5 germline | P-1 L4 | 21.5 +/- 0.5959 | 22 | <0.0001 | 80/90 | 5B |
| *dimt-1::AID* | JDW221 mex-5 germline | egg | 19.966 +/- 0.5115 | 20 | <0.0001 | 89/93 | 5B |
| *dimt-1::AID* | JDW221 mex-5 germline | egg->y.a. | 14.195 +/- 0.2814 | 14 | 0.9570 | 83/94 | 5B |
| *dimt-1::AID* | JDW221 mex-5 germline | y.a. | 19.519 +/- 0.4576 | 22 | <0.0001 | 75/87 | 5B |
| *dimt-1::AID* | CA1200 eft-3 ubiquitous | | 14.152 +/- 0.3988 | 14 | | 67/90 | 5C |
| *dimt-1::AID* | CA1200 eft-3 ubiquitous | P-1 L4 | 20.159 +/- 0.5473 | 22 | <0.0001 | 69/93 | 5C |
| *dimt-1::AID* | CA1200 eft-3 ubiquitous | egg | 18.464 +/- 0.6297 | 20 | <0.0001 | 69/90 | 5C |
| *dimt-1::AID* | CA1200 eft-3 ubiquitous | egg->y.a. | 15.588 +/- 0.5234 | 16 | 0.0101 | 68/91 | 5C |
| *dimt-1::AID* | CA1200 eft-3 ubiquitous | y.a. | 19.169 +/- 0.6477 | 22 | <0.0001 | 66/95 | 5C |
| *dimt-1::AID* | JDW221 mex-5 germline | | 13.96 +/- 0.4276 | 14 | | 53/94 | |
| *dimt-1::AID* | JDW221 mex-5 germline | P-1 L4 | 21.74 +/- 0.9826 | 19 | <0.0001 | 68/91 | |
| *dimt-1::AID* | JDW221 mex-5 germline | egg | 20.78 +/- 0.7348 | 21 | <0.0001 | 85/94 | |
| *dimt-1::AID* | JDW221 mex-5 germline | egg->y.a. | 15.21 +/- 0.4364 | 14 | 0.1426 | 66/80 | |
| *dimt-1::AID* | JDW221 mex-5 germline | y.a. | 19.69 +/- 0.8134 | 21 | <0.0001 | 64/90 | |
| *dimt-1::AID* | CA1200 eft-3 ubiquitous | | 13.93 +/- 0.5767 | 14 | | 42/87 | |
| *dimt-1::AID* | CA1200 eft-3 ubiquitous | P-1 L4 | 17.79 +/- 0.7352 | 19 | <0.0001 | 61/101 | |
| *dimt-1::AID* | CA1200 eft-3 ubiquitous | egg | 17.82 +/- 0.8017 | 19 | <0.0001 | 65/96 | |
| *dimt-1::AID* | CA1200 eft-3 ubiquitous | egg->y.a. | 15.35 +/- 0.7266 | 16 | 0.0890 | 63/96 | |
| *dimt-1::AID* | CA1200 eft-3 ubiquitous | y.a. | 17.18 +/- 0.7952 | 16 | 0.0004 | 55/92 | |
| *dimt-1::AID* | JDW221 mex-5 germline | | 18.24 +/- 0.4745 | 17 | | 75/90 | |
| *dimt-1::AID* | JDW221 mex-5 germline | y.a. | 21.182 +/- 0.5970 | 22 | <0.0001 | 77/90 | |
| *dimt-1::AID* | JDW221 mex-5 germline | PEL | 22.829 +/- 0.6534 | 22 | <0.0001 | 70/90 | |
| *dimt-1::AID* | JDW221 mex-5 germline | ML | 21.227 +/- 0.6159 | 22 | 0.0002 | 66/90 | |
| *dimt-1::AID* | JDW221 mex-5 germline | | 18.136 +/- 0.4999 | 19 | | 81/91 | 5D |
| *dimt-1::AID* | JDW221 mex-5 germline | y.a. | 22.056 +/- 0.6425 | 21 | <0.0001 | 89/91 | 5D |
| *dimt-1::AID* | JDW221 mex-5 germline | PEL | 22.671 +/- 0.6702 | 23 | <0.0001 | 70/90 | 5D |
| *dimt-1::AID* | JDW221 mex-5 germline | ML | 21.569 +/- 0.6783 | 21 | <0.0001 | 65/90 | 5D |

**Supplementary Table 5. DIMT-1 functions after mid-life to regulate lifespan** The figure panels in which specific experiments are shown or used are indicated in the right column. The mean lifespan and SD values were calculated by Prism from triplicate samples of 30 worms each (90 worms total). # worms: number of observed dead worms at the end of the experiment/number of alive worms at the beginning of the experiment. The difference between both numbers corresponds to the number of censored worms (worms that underwent “matricide”, exhibited ruptured vulva, or crawled off the plates). P values are calculated by log rank (Mantel-Cox) statistical test.
